# Micro RNA mediated enhanced growth in *Solanum lycopersicum* during cadmium hormesis

**DOI:** 10.1101/2025.07.04.663119

**Authors:** Risha Ravi, Manish Yadav, Santosh Kanade

## Abstract

Hormesis refers to the adaptive mechanism of organisms in response to environmental challenges where a lower dose of a toxic compound induce an improvement in functionality and overall development and a higher dose endangers even the existence of the organism. The recent developments in hormetic studies are of paramount importance in plant research as they help in risk assessment of environmental contaminants, protect the vegetation against pollution, and improve crop productivity. As one of the most toxic contaminants, cadmium is considered to have detrimental effects on the growth and development of plants. However, recent studies have revealed the beneficial effects of cadmium in plants at low levels of exposure however the exact mechanism behind this phenomenon is poorly deciphered. In this study, we have focused on observing the response of tomato seedlings under different concentrations of cadmium. The morphological, biochemical, and histochemical characterization of these seedlings under low cadmium exposure has confirmed their hormetic effects. The differential gene expression by transcriptomic profiling in low cadmium showed that, apart from genes involved in oxidoreductase activity, and signaling, several lncRNAs, also differentially expressed. The lncRNAs are known to regulate gene expression on the chromatin level and post-transcriptional regulation. First time we are reporting the expression of lncRNAs in hormesis as important factor for enhanced growth. In-silico analysis revealed the functions of lncRNAs, involving the prediction of cis-targets, mi-RNA precursors, and their targets. Two miRNAs; sly-MIR396a and sly-MIR1063g were seen to have a direct role in improving the growth of plants treated with low cadmium provided an insight into the molecular mechanisms of their role in cadmium hormesis. These findings provided important understanding of the molecular basis of the hormetic phenomenon which can pave a path for generating crops with improved agronomic characteristics.

**Graphical abstract:** 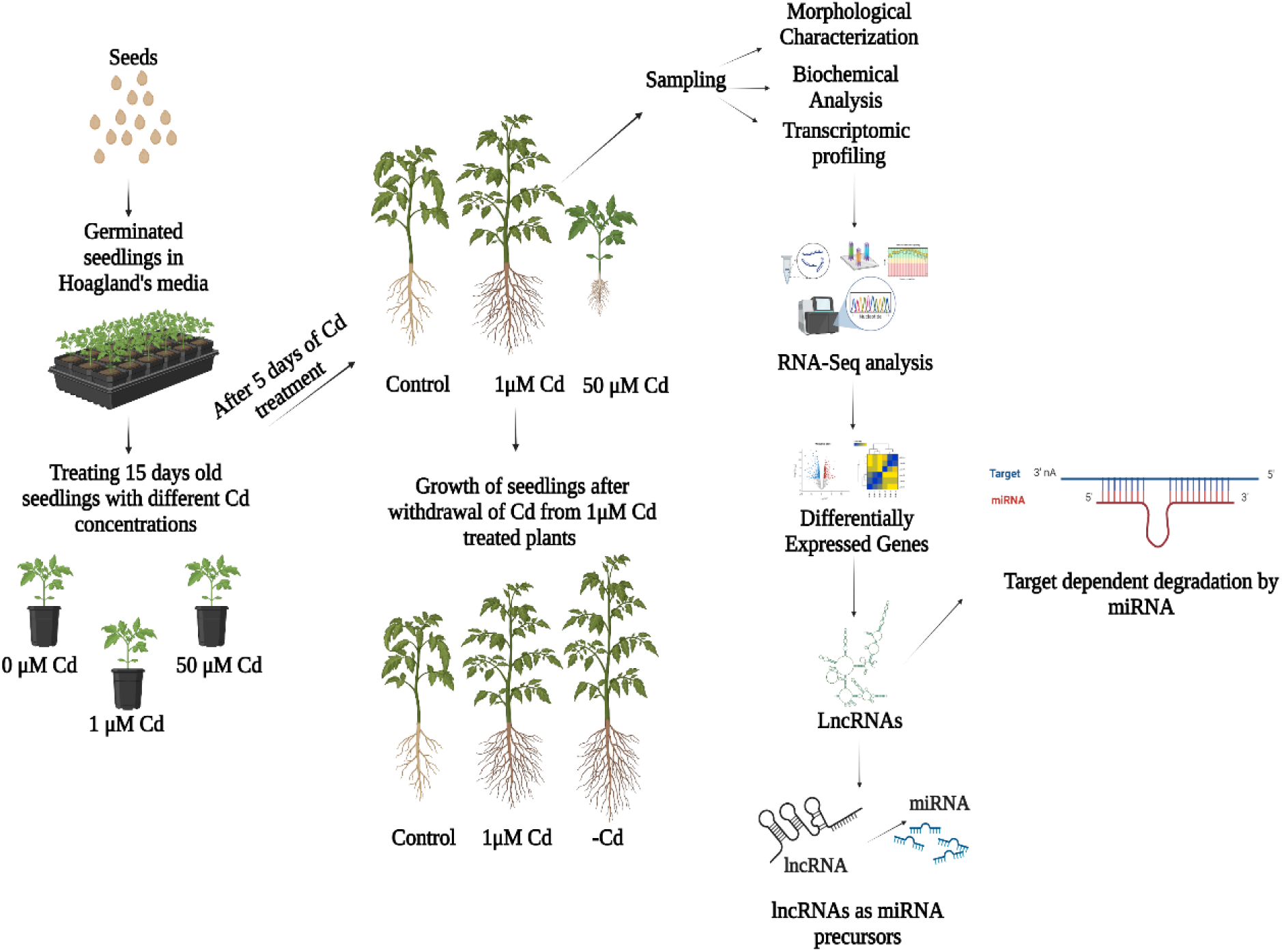

**Highlights:**

- Low cadmium exposure on *Solanum lycopersicum* seedlings showed more promising outcomes in terms of growth and development.
- The plants exposed to 1µM Cd treatment continued to exhibit superior growth responses despite removing the cadmium from the media after 5 days of treatment.
- The GO analysis of Differentially Expressed Genes (DEGs) suggested several lncRNAs differentially expressed in 1 µM Cd condition.
- Differentially expressed LncRNAs Solyc01g006780.4 and Solyc12g019150.1 generated the miRNAs, sly-MIR396a and sly-MIR1063g respectively.
- Upregulation of sly-MIR396a and sly-MIR1063g resulted in the downregulation of GRF12 and NET4B-like proteins respectively leading to increased growth of tomato plants.

## Introduction

Heavy metals like Cadmium (Cd) are known to be hazardous and a major pollutant in farmlands. According to the US Environmental Protection Agency (EPA), cadmium ranks as the third most hazardous environmental contaminant, and type III Carcinogen following mercury and lead. It is also one of the main factors that limit crop quality and productivity (Dal Corso et al., 2008; Satarug et al., 2009). Nevertheless, at low concentrations, several toxic chemical stressors benefit plant growth and development (Agathokleous et al., 2020), and this biphasic dose-response phenomenon is called hormesis. A reverse U-shaped biphasic curve is a defining feature of this phenomenon, showing a stimulatory effect in low concentrations of toxic substances. However, the lethal effect can be observed after increasing the concentrations. This is a universal phenomenon and widespread in nature, independent of the stressor type, the organism in which it occurs, or the physiological processes (Carvalho et al., 2020). Previous studies indicate that the dose-response relationship functions as a biological compass, guiding organisms through adversity and enabling adaptation for enhanced survival, particularly in response to environmental factors that disrupt the organism’s homeostasis (Calabrese & Mattson, 2017; Agathokleous et al., 2018).

Hormesis is a complex process that has not yet received enough attention. Pb and Cd showed distinct physicochemical characteristics on the hormetic stimulation of maize shoots (Małkowski et al., 2020). The occurrence of hormetic phenomenon ranges from prokaryotes to eukaryotes. With regard to bacterial studies, Iavicoli et al., 2021, explain how hormesis due to antibiotics significantly influences bacterial behaviour, helping them to endure in upcoming harsh environments. The progressive work by Shanmugam et al., 2018, suggests that the stress response in *C. elegans* may be triggered by hormetic dietary phytochemicals found in *(Dioscorea spp.)* tubers via the SKN-1/Nrf2 and HSF-1 signaling pathways. In the modern growing era, much work has been done showing how pharmaceutical products exhibit hormetic effects (Calabrese et al., 2008a). Riluzole is the longest-approved medication for treating Amyotrophic Lateral Sclerosis (ALS), and it works proactively by inducing hormetic processes (Calabrese et al., 2008c). Similarly, the hormetic dose response is present in all Alzheimer’s disease (AD) medications that the US FDA (Food and Drug Administration) has approved for use in humans (Calabrese et al., 2008b). Another application of pharmaceutical products has been reported by Zhang et al., 2017 where a low dosage of PTS (Panaxatriol saponins) could ameliorate the behaviour movement impairment in zebrafish by decreasing the 6-OHDA (6-hydroxydopamine) -induced dopaminergic neuron loss. This study indicates that PTS can be the potential bioactive compound for the development of future therapeutic medicines for Parkinson’s Disease (PD). In addition to this, Salinitro et al., 2021, suggest that the presence of low levels of toxic metals like Cd and Chromium (Cr) in urban soils might enhance the resilience of *Poaannua L.,Cardamine hirsuta L., and Stellaria media*, enabling them to better survive in human-altered environments. Low-dose glyphosate application to waterlogged plants promoted dry mass accumulation in the roots, which may be connected to an increase in carbohydrates for energy production in low-oxygen environments. These findings of Velasco et al., 2023 suggest that lettuce plants may exhibit a potential hormetic effect because of glyphosate and may be helpful in minimizing waterlogging damage to lettuce plants.

Plants live continuously with different biotic and abiotic factors, giving rise to complex interactions that severely impact crop productivity. For instance, it has been claimed that abiotic stresses can result in production losses exceeding 50% (Calanca et al., 2017), while biotic stresses are expected to cause yield losses of approximately 35% (Savary et al., 2012). Plants use molecular, cellular, biochemical, and physiological modifications as part of their response mechanisms to various abiotic and biotic factors, where most of these reactions are governed by certain genes (Erpen et al., 2018). For example, *OsMPK5, a* member of the MAPK gene family acts as a key player in the rice stress-response network as a positive regulator of both abiotic stress tolerance and ABA-controlled resistance to brown spots (Bailey et al., 2009; Xiong and Yang, 2003). Yogendra et al. 2015, showed that the pathway of HSP17.8 has been involved in biotic interactions during the infection of *Phytophthora infestans* in *Solanum tuberosum*. Several biotic and abiotic factors also influence epigenetic regulators like DNA methylation and histone modifications inside a plant (Lämke et al., 2017). Transcription factors like *ANAC055* (Tran et al., 2004), *ZmMYBIF35* (Bailo et al., 2019), and *WRKY30* (Bailo et al., 2023) are induced by salinity, chilling, and drought stress, respectively to fight and cope against these stresses. miRNA is another important regulator in biotic and abiotic stress. Yang et al., 2017, showed that 412 sugarcane miRNA has been found to alter plant growth in response to various stress circumstances.

The two primary subgroups of RNAs involved in transcription control are coding and non-coding genomic elements, which include messenger RNA (mRNA) and long non-coding RNA (lncRNA). Generally, RNA transcripts lacking coding potential and with a minimum length of 200 are referred to as lncRNAs (Chekanova et al., 2015). LncRNAs are broadly categorized according to their location relative to protein-coding genes, encompassing long intergenic non-coding RNA (lincRNA), intronic lncRNA (incRNA), sense lncRNA, and (Laurent et al., 2015; Lina Ma et al., 2013). The ongoing advancements in high-throughput sequencing, a significant number of lncRNAs have been discovered in plants in recent years (Liu et al., 2012). At first, it was believed that lncRNAs were merely byproducts of transcription because of lacking significant evolutionary conservation. However, mounting evidence indicates that lncRNAs predominantly engage with DNA, mRNA, protein, and miRNA, thereby playing essential roles across epigenetic, transcriptional, post-transcriptional, translational, and posttranslational dimensions (Kung et al., 2013; Lucero et al., 2021) modulating diverse biological functions, such as plant growth, development, and responses to environmental stresses (Rinn et al., 2012; Chekanova et al., 2015; Liu et al., 2012, and Sun et al., 2019). LncRNAs additionally function as enhancer RNAs, facilitating the recruitment of RNA polymerase II to gene promoters, thus enhancing gene transcription (Kim et al., 2011). Moreover, certain lncRNAs have the ability to function as endogenous target mimics (eTMs) of miRNAs, where they can engage in competitive interactions with mRNAs for binding to miRNAs and lessen the inhibitory effect of miRNAs (Franco-Zorrilla et al., 2007).

Currently, no living system has shown any indication that a hormetic phenomenon caused by lncRNA has occurred. Even though there is ample evidence linking lncRNA to both biotic and abiotic stress in plant as well as animal kingdom. The lncRNA2 and lncRNA7 of cotton can regulate the genes involved in plant resistance to *V. dahliae* and other multiple fungal pathogens by cell wall remodeling (Lin et al., 2022). Reports by Wang et al., 2018b suggests that out of 1411 lncRNAs, 239 lncRNAs were differentially expressed in tomatoes under low-temperature stress. Furthermore, it has been found that lncRNA1459 regulates the ripening process of tomato fruits. The lncRNA1459 mutant’s fruits displayed lower levels of ethylene during the ripening stages than those of the wild type (Li et al., 2018). The expression of lncRNA (MEG3) is notably elevated in osteosarcoma cell lines, and the knockdown of MEG3 suppresses cell proliferation and metastasis while promoting apoptosis in osteosarcoma cells (Zhao et al., 2018). While numerous studies have explored stress and developmental responses in plants regarding lncRNAs, there remains a substantial need for further investigation to map lncRNAs and their potential involvement in hormesis comprehensively. In present study, plants treated with low Cd concentrations showed better growth response than the control and high Cd concentrations. The stress markers like total MDA (Malondialdehyde) and total proline content were significantly reduced than the control. Throughput transcriptome sequencing helped us to identify a few of the deferentially expressed lncRNA in the hormesis condition. We have also validated the potential targets of lncRNA bioinformatically, which was validated by RT-PCR. We hypothesize that this lncRNA might be involved in regulating the gene responsible for growth and development.

## Materials and Methods

### Materials

Arka Samrat variety of tomato seeds were procured from IIHR Bangalore India; Analytical grades of 3,3′-diaminobenzidine (DAB) (RM2735), Evans blue (GRM942), Nitro-Blue Tetrazolium (NBT) (RM578), Trichloro acetic acid (GRM7570), Thiobarbituric acid (RM1594), Sulphosalicylic acid 3% (R020), L-Proline (RM061), and Bradford reagent (ML106) was procured from HiMedia Laboratories Private Limited, India. Cadmium chloride (02417) was purchased from Loba Chemie Pvt. Ltd and Phenylmethyl sulfonyl fluoride (PMSF) (87606) was purchased from Sisco Research Laboratories Pvt. Ltd.(SRL) - India. Spectral^TM^ Plant Total RNA Kit (STRN 50, SIGMA) was obtained from Sigma. Prime Script™ 1st strand cDNA Synthesis Kit (6110A, TaKaRa) and TB Green *Premix Ex Taq* II (Tli RNase H Plus) (RR82LR, TaKaRa) were purchased from DSS Takara Bio India Pvt. Ltd.

## Methods

### Plant growth and treatment

Tomato seeds of the Arka Samrat variety, purchased from IIHR Bangalore were used for the study. Surface sterilized (2.5% sodium hypochlorite for 5 minutes) seeds were washed in running tap water 3-4 times and sown in trays with double-layered filter paper in distilled water in the dark for 5 days. Germinated seeds were transferred to pots containing half-strength Hoagland media (pH-5.8) and grown in controlled conditions at a temperature of 24^0^C, with a 16hr light / 8hr dark photoperiod at a light intensity of 200µmol/m^-2^ s^-1^. For the treatment, different concentrations (0, 0.5, 1, 2, & 50µM) of cadmium chloride solution were directly added to the media, and plants were grown for 5 more days. The media was renewed every five days to avoid fluctuations in the pH and nutrient depletion. After 20 days the plants were harvested for further experiments.

To check the residual effects of cadmium on tomato plants, after 5 days of treatment,1 µM treated plants were taken and grown for another 10 days by removing the media with normal half-strength Hoagland’s without cadmium along with control (0µM Cd), and 1µM Cd-treated plants. The basic growth parameters, such as plant length and biomass, were measured every three days during the entire treatment.

### Determination of Plant Growth

The 20-day-old plants were harvested; the height, fresh weight, and dry weight of roots and shoots were measured manually. The plant height; root and shoot length were measured separately with a ruler. The fresh weight of the seedlings was determined using a weighing balance after dabbing with filter paper to soak the extra media from root surfaces. To calculate the dry weight, plant samples were dried at 60^0^C in an oven up to constant weight and weighed.

### MDA Content estimation

The MDA content from roots and leaves was estimated according to the thiobarbituric acid (TBA) method with slight modifications (Meng et al., 2022). Plant samples were powdered in liquid nitrogen and 100 mg were homogenized in 1ml of 5% Trichloro acetic acid (TCA) solution containing 0.25% TBA. The homogenate was boiled at 100°C in the water bath for about 30 minutes and centrifuged at 13,000 x g for 15 minutes, after being cooled to room temperature. The absorbance of the supernatant was measured at 440, 532, and 600 nm using Spark® multimode microplate reader (Tecan Spark Control magellan), and MDA content was calculated using the formula: MDA content = [6.45 × (A532 - A600) - 0.56 × A440] × V/FW.

### Estimation of Proline content

Total proline content in leaves and roots of control and treated plants was estimated using the method (Bates et al., 1973). Plant samples were ground to fine powder using liquid nitrogen and 100 mg of powder was homogenized in 0.5ml of 3% sulfosalicylic acid. The homogenate was centrifuged at 12,000xg for 15 minutes. An aliquot of the supernatant (100µl) was added to 200µl of acid-ninhydrin and 200µl of glacial acetic acid and boiled at 100^0^C for one hour. The reaction was terminated by keeping them in ice. One ml of toluene was added to the reaction mixture, and incubated at room temperature for about 10 minutes after vigorous stirring. The toluene layer containing the proline-ninhydrin chromophore was collected and absorbance was measured at 520 nm using toluene as a blank in Spark® multimode microplate reader (Tecan Spark Control magellan). The proline content in µg/ml was obtained from the standard graph for proline and the corresponding proline in µmoles/g FW was calculated using the formula: Proline (µmoles/g FW) = µg/ml proline x ml toluene /115.5 x 5/g sample.

### Relative Leaf Water Content

The relative leaf water content of leaves was measured according to the method described by Yamasaki and Dillenburg (1999). Leaves of control and treated plants were collected and fresh weight was measured. The leaves were allowed to be floated on distilled water for one hour to record the turgid weight. They were oven-dried for 24 hours and dry weight were measured. RLWC was calculated using the formula: RLWC% = [Fresh weight - Dry weight / Turgid weight - Dry weight] x 100.

### Anti-oxidant enzyme assays

The leaf samples (100mg) were ground in liquid nitrogen and homogenized in 50 mM phosphate buffer, pH 7.0, containing 1 mM PMSF. The samples were centrifuged at 10,000 rpm for 10 minutes at 4^0^ °C, and the supernatant was taken. The protein concentration was estimated using Bradford method and used for catalase and SOD assays.

### Catalase estimation

The catalase activity was measured using the method of Patterson et al. (1984). The reaction mixture contained 50 mM phosphate buffer (pH 7.0), 20 mM H_2_O_2,_ and 25 µg protein made up to 1ml. The oxidation of H_2_O_2_ was measured spectrophotometrically at 240 nm, and the catalase activity was measured accordingly using the extinction coefficient of H_2_O_2_ at 240 nm (43.6 mM-1 cm-1).

### Superoxide dismutase enzyme activity

The SOD activity was estimated according to the method devised by Beauchamp and Fridovich (1971). The reaction mixture consisted of 27 ml of sodium phosphate buffer (pH 7.8), 1 ml of NBT (1.44 mg ml-1), 1.5 ml of methionine (30mg ml-1), 1.5 ml of 2 mM EDTA, 0.75 ml of Triton X-100. 10 µl of riboflavin (4.4 mg 100 ml-1) and enzyme extract containing 50 µg of protein were added to 1 ml of this reaction mixture and illuminated for 8 minutes under three comptalaux bulbs at 25 o C. A tube without extract but kept in light would serve as a control, and a tube with extract kept in the dark serves as the blank. The difference in the NBT reduction at 560 nm in light with and without enzyme is measured as the SOD activity. The amount of protein required to inhibit 50% initial reduction of NBT under light is considered as one unit of enzyme activity.

### Histochemical Analysis

#### Qualitative estimation of Superoxide Anions

Superoxide anion radicals were qualitatively estimated and visualised by NBT staining (Kumar et al., 2014). The 0.2% NBT staining solution was prepared by dissolving 0.25g NBT in 50mM sodium phosphate buffer (pH 7.5). Two to three leaves each from control and treatments were harvested and immersed in 0.05% NBT solution at room temperature in a shaker for 4-5 hrs at 60-80 rpm. The leaves were transferred to a bleaching solution (Ethanol: Glycerol: Acetic acid (3:1:1)) and boiled at 95°C in a water bath for 30 minutes to terminate the reaction and to remove the chlorophyll pigments. The dark blue spots on the decolourized leaves indicating the superoxide radicals were then visualized and photographed.

#### Qualitative estimation of Hydrogen peroxide

Qualitative estimation of hydrogen peroxide radicals was performed by DAB staining described by Daudi and O’Brien (2012). Leaves were immersed in DAB solution (1mg/ml) and incubated for 4 hrs in the dark. The staining solution was replaced with bleaching solution (Ethanol: Glycerol: Acetic acid (3:1:1)) and boiled at 95°C in a water bath for 30 minutes. The leaves were further immersed in fresh bleaching solution for 30 minutes and photographed.

#### Evans blue staining

The damage to the root of the seedlings was assessed by Evans blue staining (Baker and Mock, 1994). After immersing the roots in Evans blue solution (0.25%, w/v) in a shaker at room temperature for 30 minutes, they were destained using distilled water. Subsequently, the samples were visualized in a light microscope and photographed.

### RNA isolation and Transcriptomic analysis

RNA was isolated from the root and leaves of control and treated plant samples using a Spectral plant total RNA kit (STRN50, Sigma) according to the manufacturer instructions and given for RNA-Seq analysis at Novel ge ne Technologies Pvt Ltd, Hyderabad, India. RNA concentration and integrity were checked using a Nanodrop *(NanoDrop™2000/2000c Spectrophotometer,* Thermo Scientific™) and bioanalyzer (Agilent 2100 Bioanalyzer, Agilent Technologies) respectively. The purity and contamination of RNA were detected using a 1% agarose gel. Purified and intact RNA were used to prepare a cDNA-library and the high-quality reads were sequenced on Illumina Nova. Seq6000.

#### Mapping and data filtering

The raw data obtained from the sequencer were processed by removing adapter sequences and selecting high-quality reads on the parameter of having>Q20 value. All the high-quality reads were mapped against the reference *Solanum lycopersicum* (tomato:-GCF000188115.4) genome using the Hisat2 genome assembler.

#### Differential gene expression and GO Enrichment analysis

The annotation of the transcriptome count table was created using the Feature counts option of the R subread package using the gene transfer format (GTF) as a reference. The assembled transcriptome expression was estimated using the DESeq2package of R. The Majority of the assembled transcripts have expression>=1FPKM. Gene Ontology (GO) annotations for the DEGs were predicted using Blast2GO software. The significantly enriched GO terms and pathways were selected for analysis.

#### Identification of lncRNAs

Uncharacterized transcripts longer than 200 nucleotides were selected and uploaded to the InterPro software (https://www.ebi.ac.uk/interpro/) to check whether they belong to any protein families. The selected sequences were later searched for ORFs using NCBI OFR Finder (https://www.ncbi.nlm.nih.gov › orffinder) and candidate lncRNAs were screened for their coding potential using *Coding Potential Calculator 2.0* (http://cpc2.cbi.pku.edu.cn) and Plant ncRNA database (PNRD) (http://structuralbiology.cau.edu.cn/PNRD), and common non-coding candidates were further studied.

#### lncRNAs as miRNA precursors

To predict the probable precursors of miRNA, lncRNAs were blasted against the miRBase (http://www.mirbase.org/search.shtml), and the identity between lncRNAs and miRNA precursors that were 100 % and e-value < 0.001 were selected. The predicted miRNA precursor lncRNAs were further analyzed for their secondary structure using the Vienna RNA fold web server (http://rna.tbi.univie.ac.at).

#### Identification of miRNA targets

The miRNA targets were predicted using the psRNATarget software and compared with our transcriptomic data. The interaction between the miRNAs and the mRNAs was analyzed using default parameters of a maximum expectation of 2.0. DEGs with significant fold change were selected and analyzed for their functions.

#### Phylogenetic and structural analysis of GRFs in tomato

The GRF sequences of tomato, as well as other relevant GRFs from maize, poplar, and rice, were retrieved and aligned using ClustalW. A Neighbor-joining phylogenetic tree was generated using MEGA12 software (https://www.megasoftware.net). The method used 1000 bootstrap values, a Poisson model with uniform rates, and pairwise deletion. The MEME Suite database (https://meme-suite.org) was used for checking the structural properties and conserved domains of the GRF genes.

#### Interaction studies between miRNAs and Targets

The RNAhybrid (https://bibiserv.cebitec.uni-bielefeld.de ›rnahybrid) tool was used to check the relationship between miRNAs and their targets. The tool produces the minimum free energy hybridization, which predicts the interaction between miRNAs and their targets.

#### Phenotypic analysis of leaves

Leaf phenotypic analysis was performed according to the method of Wang et al., 2020. The third completely developed leaf from control and treatments was taken and fixed in FAA solution (formaldehyde: acetic acid: 96% alcohol: water; 10:5:50:35) overnight at 4 °C and cleared with chloral solution (200 g chloral hydrate, 20 g glycerol, and 50 mL dH_2_O). the leaves were then observed using a Differential Interference Contrast (DIC) microscope. The epidermal cells were observed, and the cell area and cell number were calculated.

#### Q-RTPCR validation

The RNA was isolated from samples using a Spectral plant total RNA kit (STRN50, Sigma), and 1µg of RNA was taken to prepare cDNA using the PrimeScript™ 1st strand cDNA Synthesis Kit (6110 B, TaKaRa) according to the manufacturer’s instructions using the oligo dT primers. The PCR was carried out in a 10 μl reaction volume containing 1X buffer with SYBR Green (TB Green *Premix Ex Taq* II (Tli RNase H Plus) (RR82LR, TaKaRa)) and 10 μM primers. PCR was done with specific primers for selected genes (Supplementary Table 3). Ubiquitin and Actin primers were used as internal controls. Following reactions is followed for amplification initial denaturation at 95 C for 3 minutes followed by denaturation at 95 C for 10 sec annealing at 60C for 30 sec denaturation 95 C and stored at 4 C. The primer list provided Supplementary Table 3.

#### Statistical analysis

All the data were presented as mean values of each treatment. At least 2 independent biological replicates were used for each experiment. All the graphs and statistical analyses were done using GraphPad Prism 10 software. A Student’s t-test was carried out between the control and treatment groups of the same experiment to check the significance of the data obtained. P < 0.05 was considered significant. All the images were made using the BioRender software.

## RESULTS

### Influence of cadmium on the growth and development of plants

Low concentration of cadmium has an evident impact on the growth and development of plants. To study the morphological changes mediated by the hormetic effect of cadmium, we have selected the Arka Samrat variety of tomato plants. The 15-day-old tomato seedlings were treated with indicated cadmium concentrations for 5 days, and different parameters were measured. As shown in **Fig.1**, there was a significant increase in the root length, shoot length, fresh weight, and dry weight of plants exposed to lower cadmium concentrations and a notable decrease in these parameters in higher cadmium concentrations when compared to the control. The highest root length was observed in 1 µM Cd with an increase of 18.8% when compared to the control and the lowest root and shoot length was found to be in 50 µM with a decrease of 28.4% and 31.11% respectively compared to the control (Figure 1A and B). Significant enhancement was noticed in fresh weight and dry weight with 41.9% and 38.2% in 1 µM Cd whereas reduction by 61.47% and 43.4% in 50 µM respectively compared to control (Figure 1 E and F). The root architecture was presented in figure 1G with and without cadmium; results demonstrated that branching of root is significantly increased at low concentration of cadmium.

**Fig. 1:**
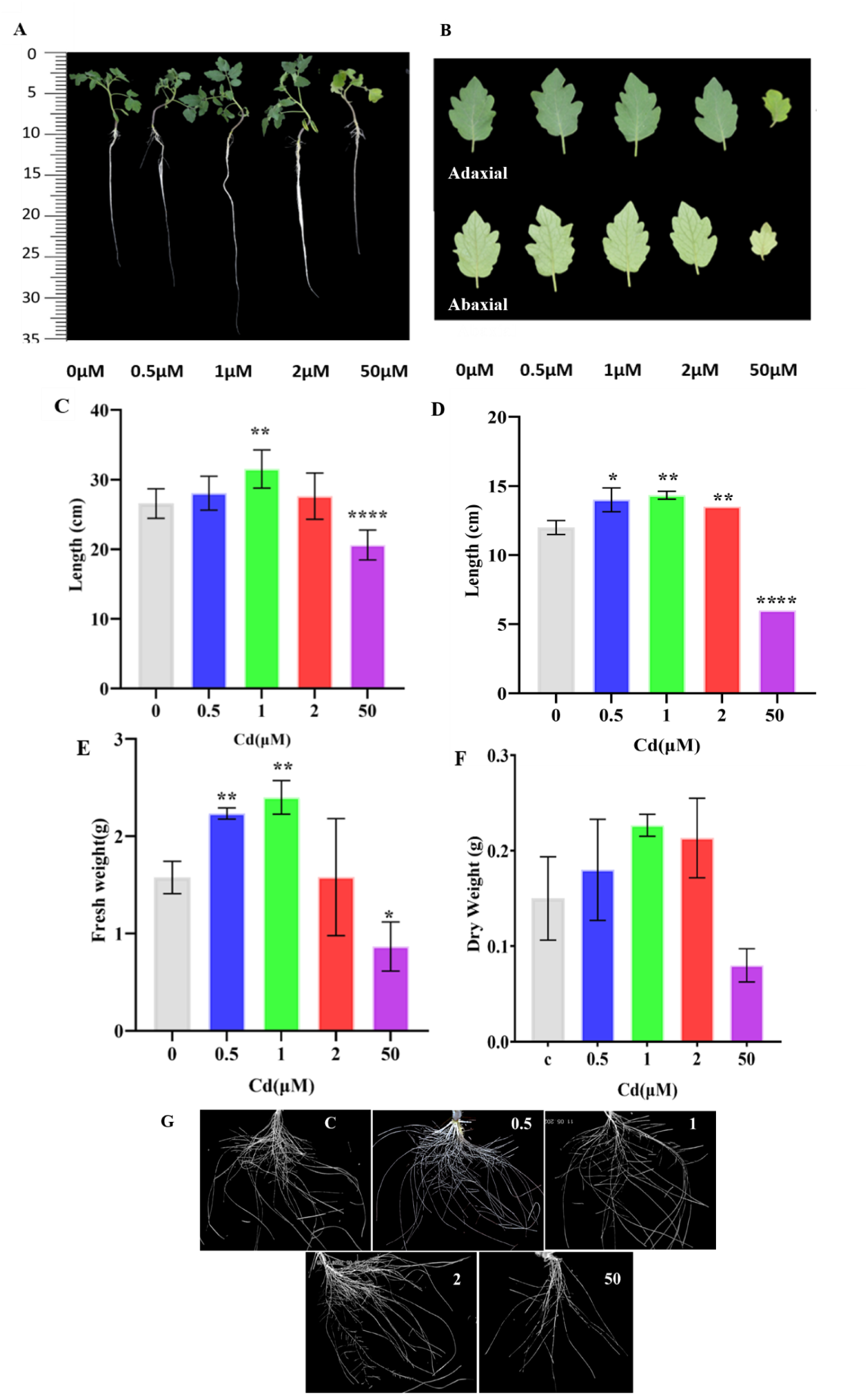
Influence of cadmium on plant growth. A) Influence of cadmium on the plant growth and B) leaf architecture, C) root length, and D) shoot length of tomato plants treated with different cadmium concentrations. Influence of cadmium on the E) Fresh weight, F) Dry weight, and root architecture of tomato plants under varying cadmium concentrations. Plant length and biomass were measured after treating 15-day-old plants with different cadmium concentrations for five days. Bars in the graph show mean ± SD calculated from 3 biological replicates. The asterisks above the graph represent statistically significant differences between the control and treatments determined by the student’s *t*-test (*p<0.05, **p <0.01, ***p<0.0005)

### Effects of cadmium on MDA, proline and Relative Leaf Water Content

The MDA and proline content of roots and leaves of tomato seedlings after 5 days of treatment are presented in **Fig.2**. The MDA is a byproduct of lipid peroxidation and commonly considered as a marker of oxidative stress. Proline is a non-essential amino acid, which acts as an osmolyte, protects sub-cellular structures, scavenges free radicals, and also helps in maintaining cellular redox potential under various stress conditions. It was found that proline accumulates in plants under stress conditions like salinity stress, drought stress, heavy metal exposure, heat stress, etc. In the present study at 1µM concentration of cadmium, the MDA and proline contents of leaves are lesser than the control, whereas in 50µM increased. Similarly in the case of roots, MDA content in 1µM concentration was comparable to the control, and proline content seemed to be increased. In 50µM, MDA content was higher and proline content was comparable to 1µM. Similar to these observations, the blue signal from plant roots subjected to Evan’s blue staining (Supplementary Fig. 2) was much less intense in control and lower cadmium treated plants compared to 50µM This clearly indicates that low cadmium concentrations are not causing significant damage to membranes compared to control and stress conditions. Leaf relative water content (RLWC) is an important indicator of water status in plants as it reflects the balance between water supply to the leaf tissue and transpiration rate. As depicted in **Fig.2 E**, the relative water content showed a 5.4% increase in plants with 1μM cadmium after 5 days of treatment whereas at the increased concentration (50μM), the relative water content in the leaves was decreased by 9.4% compared to the control. The residual effect of cadmium on tomato plants, was accessed by growing the plants without Cd after initial treatment, data suggest that as shown in the Fig 3, Cd induced effect is not decreased in terms of basic growth parameters, like plant length and biomass in experimental setup when compared to untreated and continues treatment. **RNA-Seq analysis using Illumina sequencing:** To obtain a differential gene expression at hormetic level of cadmium, transcriptomic profiling was performed in the leaves and roots of cadmium-treated and control plants with three biological replicates. An average of ∼10.1 GB data in leaves and ∼ 7.7 GB data in roots was obtained using the Illumina NovaSeq 6000 platform. After filtering, there were 63.2-82.3 MB total reads in leaves and 24.8.-56.6 MB reads in roots which had Q20 value >98% and Q30 value >94%. All the high-quality reads were mapped against the reference *Solanum lycopersicum* (ITAG4.0) genome using the Hisat 2 genome assembler where more than 98% were aligned to the genome. The transcriptome data was submitted in the NCBI database with the accession number **PRJNA1270784**.

**Fig. 2:**
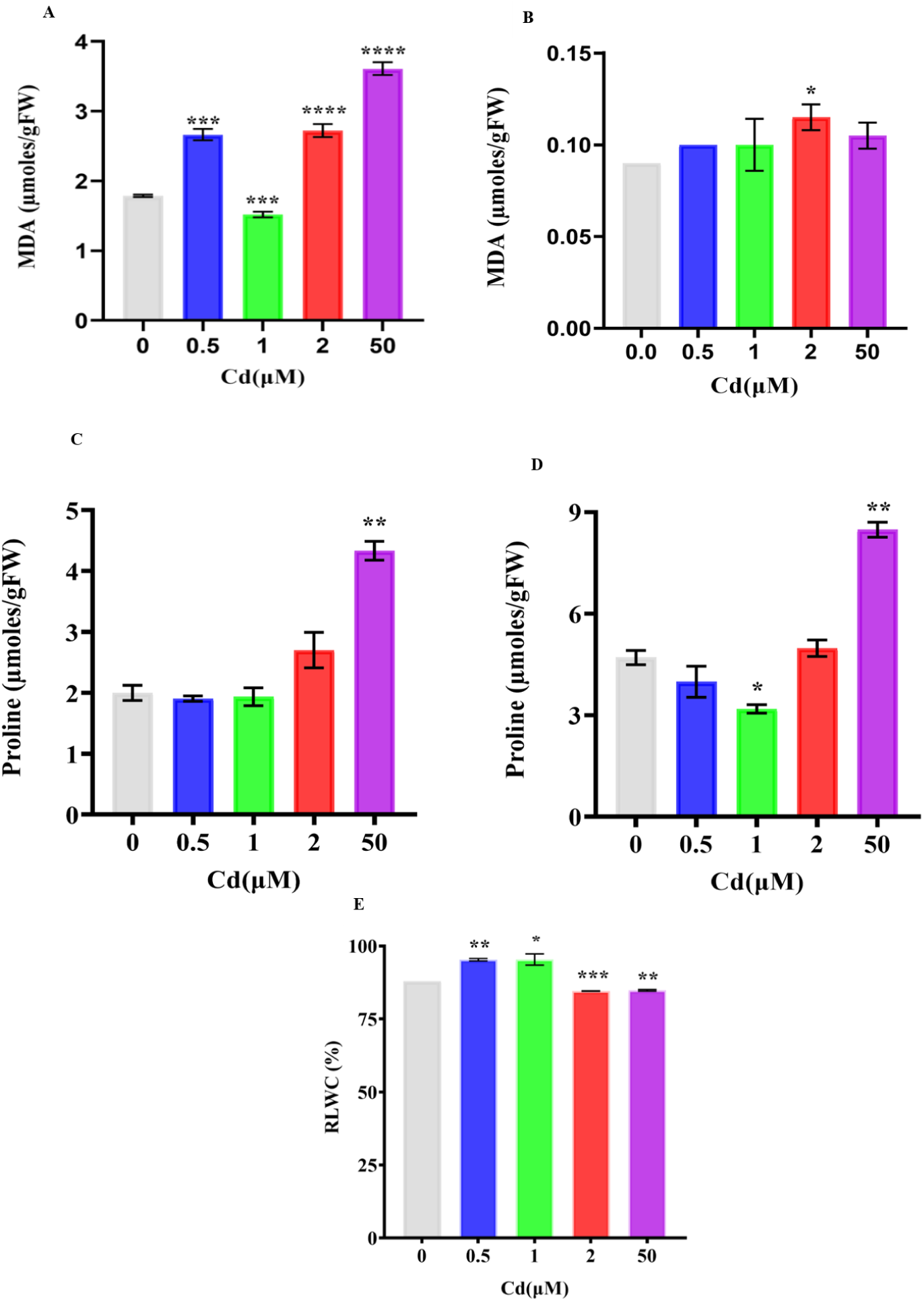
Effect of cadmium on MDA content, Proline content, and RLWC. Effect of cadmium on the MDA content of A) leaves and B) roots of tomato plants treated with different cadmium concentrations. Effect of cadmium on the Proline content of C) leaves and D) roots of tomato plants treated with different cadmium concentrations. E) RLWC of tomato plants treated with different cadmium concentrations. Bars in the graph show mean ± SD calculated from 3 biological replicates. The asterisks above the graph represent statistically significant differences between the control and treatments determined by the student’s *t*-test (*p<0.05, **p <0.01, ***p<0.0005)

**Fig. 3:**
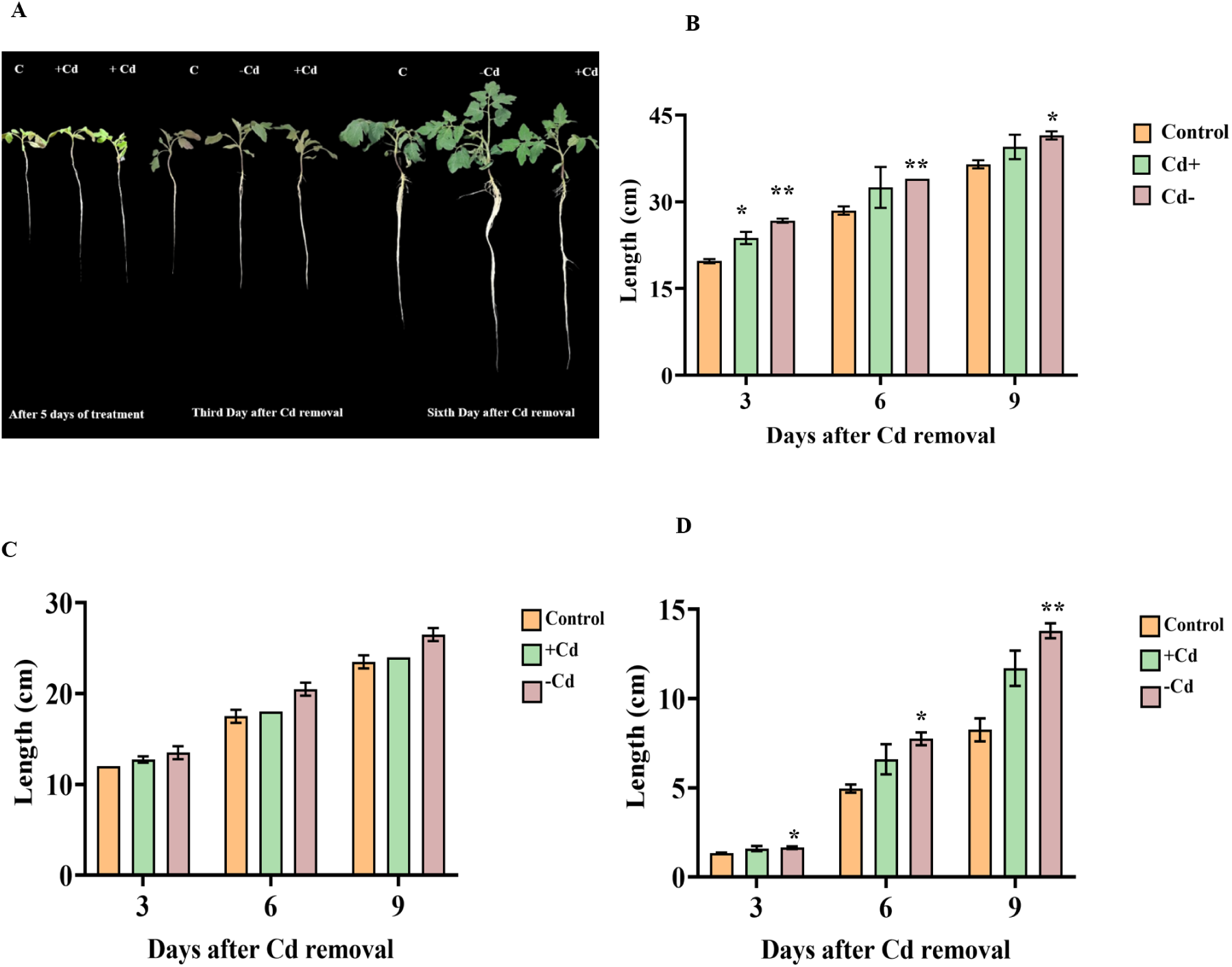
Effect on Phenotypical traits after removing cadmium from media: A) Morphological characters of tomato seedlings grown for one month after removing cadmium from the media. B) Root length, C) Shoot length and D) Biomass of tomato plants grown in cadmium-free media measured every three days for one month. After 5 days of treatment with 1µM Cd, the plants were grown in cadmium-free media for one month and the phenotypical parameters were measured every three days. The results were compared with the control and another set of tomato plants grown in 1µM Cd for one month. Bars in the graph show mean ± SD calculated from 3 biological replicates. The asterisks above the graph represent statistically significant differences between the control and treatments determined by the student’s *t*-test (*p<0.05, **p <0.01, ***p<0.0005).

### Differentially Expressed Genes in response to cadmium treatment

Differentially expressed genes in the low and high-cadmium-treated samples compared with the control were screened from the data according to the criteria of having a log fold change > 1 and p-value being ≤ 0.05 **(Fig4. A-D).** A total of 1385 genes were differentially expressed in low cadmium conditions in leaves with 599 upregulated and 786 down regulated genes. In the case of root, 507 genes were differentially expressed in which 187 genes were upregulated and 315 genes were down regulated. Compared to the low cadmium condition, the high cadmium condition relatively had a greater number of differentially expressed genes. In leaves, a total of 5932 differentially expressed genes were listed of which 3130 genes were upregulated and 2802 genes were down regulated. About 1574 genes of which 306 upregulated and 1266 down regulated genes were differentially expressed in roots under high cadmium condition. The number of unique and common genes expressed in different conditions is depicted in the form of a Venn diagram in **Fig. 4(E-F)**.

**Fig. 4:**
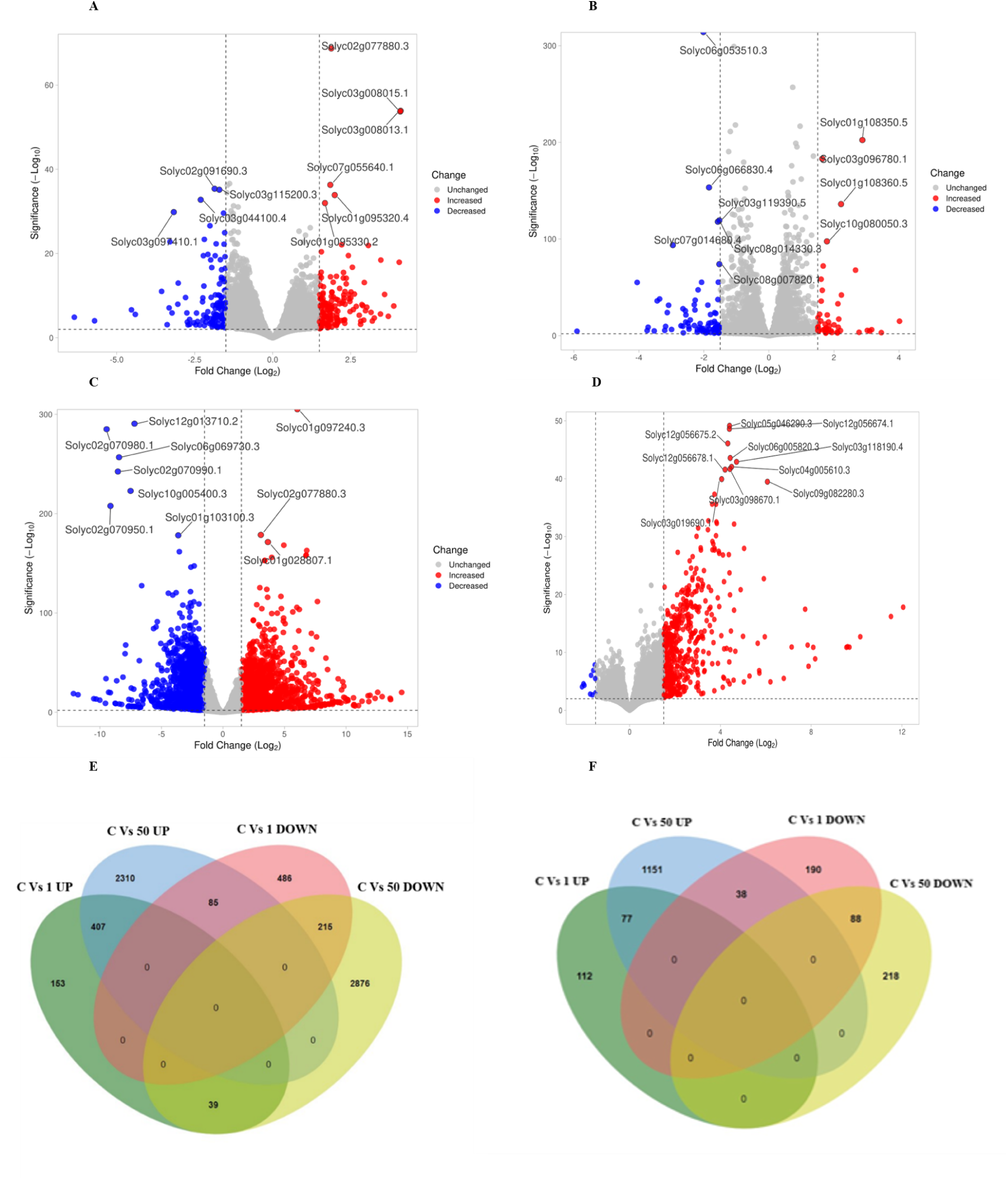
Volcano plots displaying DEGs in low and high cadmium treatments. Volcano plot showing DEGs in A) C Vs 1µM Cd, B) C Vs 50 µM Cd in leaves and C) C Vs 1µM Cd, D) C Vs 50 µM Cd in roots. Venn diagram displaying the common and unique DEGs identified in different comparisons in E) leaves and F) roots. The genes displayed in the plots have a log2(fold change) cutoff of 1 and a p-value ≤ 0.05.

### GO analysis of DEGs

To get a more comprehensive idea about the DEGs’ functions, GO analysis was performed using Blast2GO software **(Fig. 5(A-H))**. Up-regulated DEGs in leaves of low cadmium conditions were mainly enriched in transmembrane transport and phosphorylation in biological processes indicating the improved translocation of mineral nutrients and ion transport within the plant cells. In cellular components, the chloroplast and nucleus were primarily enriched which rationalizes the increased photosynthetic efficiency observed. ‘Metal ion binding’, ‘hydrolase activity’, and ‘protein binding’ are mainly enriched in the molecular function category which is related to metal chelating, signaling, and regulatory activities. In the case of down regulated DEGs, cell cycle and defensive responses in biological processes; cytoplasm, protein-containing complex, and nucleus in cellular component; transferase, hydrolase, and DNA-binding in molecular function category were enriched. In high cadmium conditions, up-regulated DEGs associated with transport and defense were enriched in the biological process category; cytoplasm and nucleus in cellular component; and metal binding and hydrolase activity were enriched in the molecular function category. DEGs related to ‘protein metabolic process’, ‘transport’, ‘gene expression, and RNA metabolomic process in the biological process; chloroplast and thylakoid in cellular components; transferase, metal ion binding, oxidoreductase activity, RNA, and protein binding in molecular function category were enriched in down regulated DEGs in high cadmium condition which explains the growth and development retardation and decreased pigments and photosynthetic efficiency observed in these plants.

**Fig. 5:**
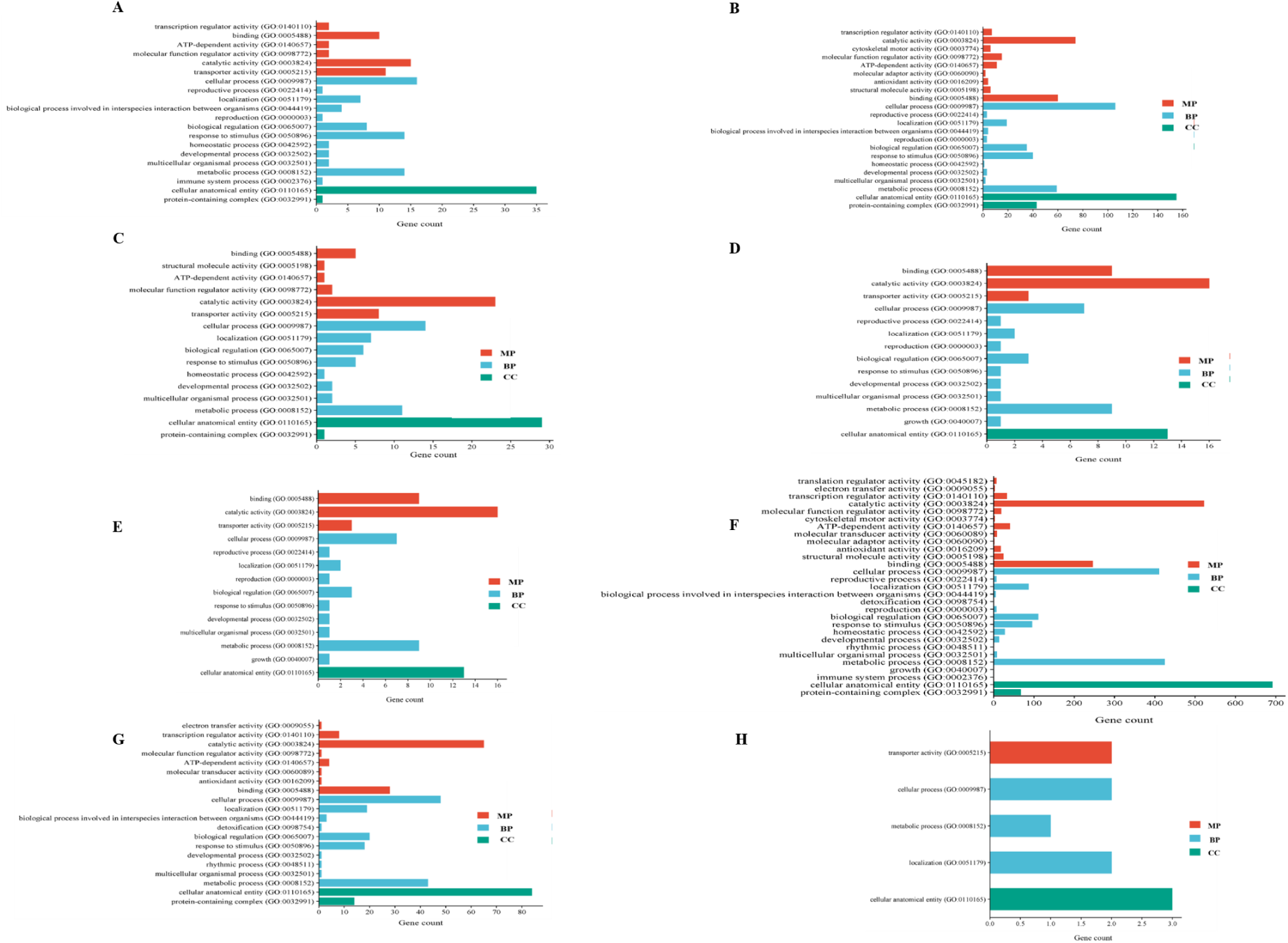
(A-D) GO Annotations of DEGs in plants treated with 1µMcadmium. GO Annotations of A) upregulated DEGs in 1µM Cd treated leaves B) downregulated DEGs in 1µM Cd leaves C) upregulated DEGs in 1µM Cd roots D) downregulated in 1µM Cd roots. **(E-H) GO Annotations of DEGs in plants treated with 50µM cadmium.** GO Annotations of E) upregulated DEGs in 50µM Cd leaves F) downregulated DEGs in 50µM leaves G) upregulated DEGs in 50µM Cd roots H) downregulated in 50µM Cd roots.

In the case of root, a similar pattern was observed where in low cadmium condition; upregulated DEGs related to transmembrane transport, transcription, and defense were predominantly enriched in biological process; cytoplasm, nucleus, and plasma membrane in cellular components and oxidoreductase activity, DNA and nucleotide binding activities were notably enriched in molecular function category. In downregulated DEGs, interestingly, defense response to fungus, phosphorylation, and hormone-mediated signaling were significantly enriched in the biological process; plasma membrane, extracellular region, and nucleus were mainly enriched in cellular component; and hydrolase activity along with protein binding were remarkably enriched in molecular function. After analyzing the DEGs in high cadmium, the upregulated DEGS contrary to the leaves were particularly enriched in transcription and phosphorylation in the biological process category; cytoplasm and nucleus in cellular components; and oxidoreductase, hydrolase, and protein and DNA binding were markedly enriched in molecular function. Strikingly the downregulated genes were also enriched in the transcription category in biological processes; plasma membrane in cellular component and oxidoreductase activity in molecular function.

### Identification of lncRNAs

Transcripts of more than 200 nucleotides obtained from RNA-Seq analysis that do not code any particular protein or have an unknown function were used to calculate their coding potential by Coding Potential Calculator (CPC) and Plant Non-coding RNA Database (PNRD). The sequences were uploaded to the InterPro software to check whether they belong to any known protein family. Transcripts having a coding potential <1 in CPC and < 0 in PNRD were considered as long non-coding transcripts. **(Supplementary Table 2)**. They were also checked using the NCBI ORF Finder tool to find functional ORFs in the sequence (**Supplementary Table 1).** The sequences that have an ORF of more than 300 nucleotides in length were omitted, and the rest were selected for further studies. After screening with these analytical tools, the rest of the lncRNAs were analysed further for their likely function.

### *Cis-*targets of lncRNAs in low cadmium-treated plants

To determine the function of lncRNAs, the potential *cis*-targets were identified by examining the coding genes 300 bp upstream and downstream of each lncRNA. The target genes obtained were compared with the differentially expressed genes observed in our analysis and investigated further. A total of 13 lncRNA-mRNA pairs were found from 11 lncRNAs and 2 differentially expressed mRNAs **(Table 1).** The GO enrichment analysis of the target genes of lncRNAs using ShinyGO 0.76 software**(Fig.6A)** predicted diterpenoid catabolic process, actin organization, DNA integrity checkpoint signaling, stomatal movement, etc., as the significantly enriched biological processes, and Gibberellin 2-beta-dioxygenase activity, DNA topoisomerase activity, Ligand-gated ion channel activity, Potassium channel activity, Water channel activity, Tubulin binding, etc. significantly enriched in the molecular function category(**Fig.6**B). The pathway enrichment analysis using the KEGG database didn’t provide any significant pathways.

**Fig. 6:**
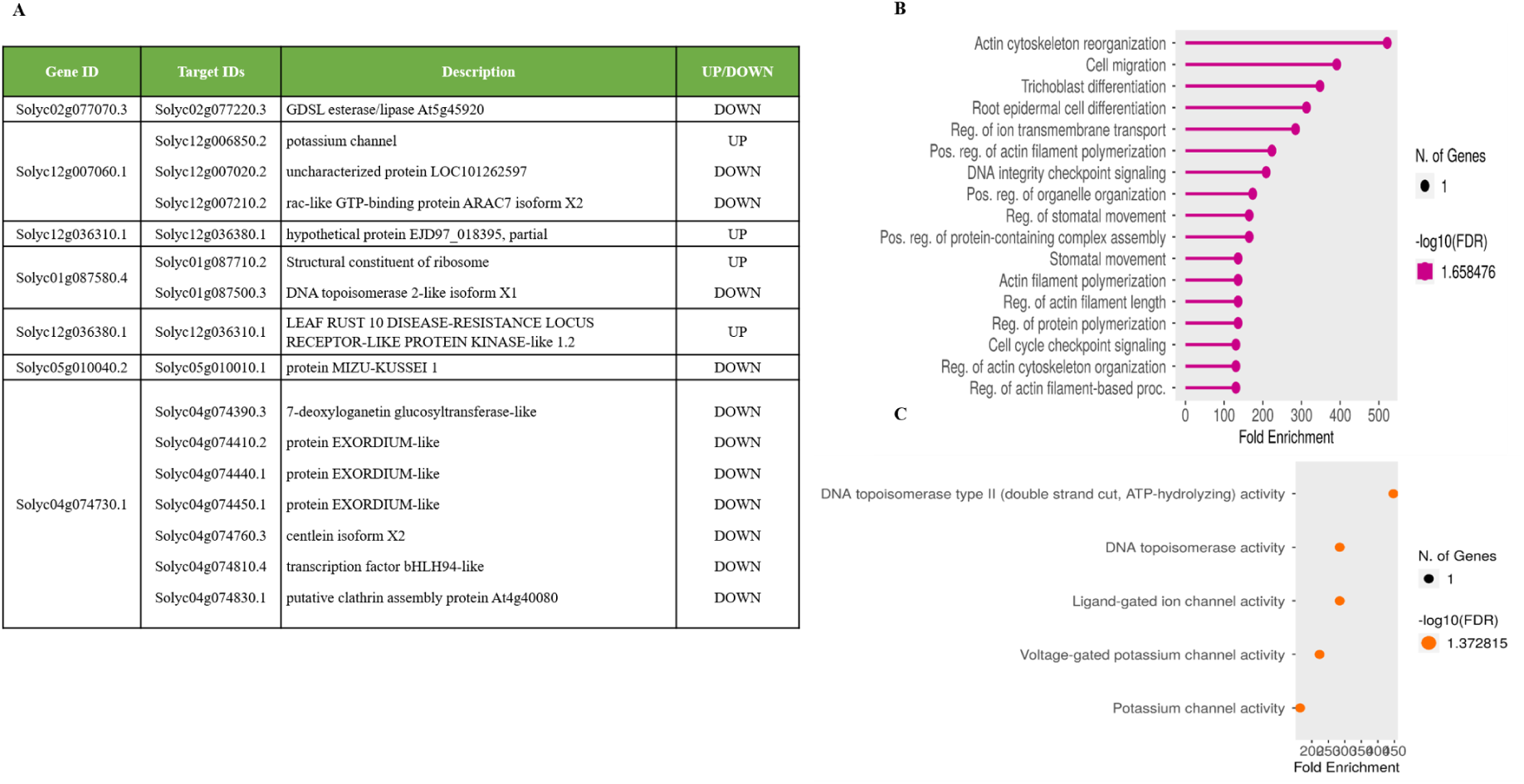
GO enrichment analysis of the cis-targets of lncRNAs using ShinyGO 0.76 software. A) Table no.1: List of LncRNAs and their predicted cis-targets. B) Dot plots showing GO enrichment analysis of molecular function and C) Biological process of differentially expressed cis-targets.

**Table no.1:**
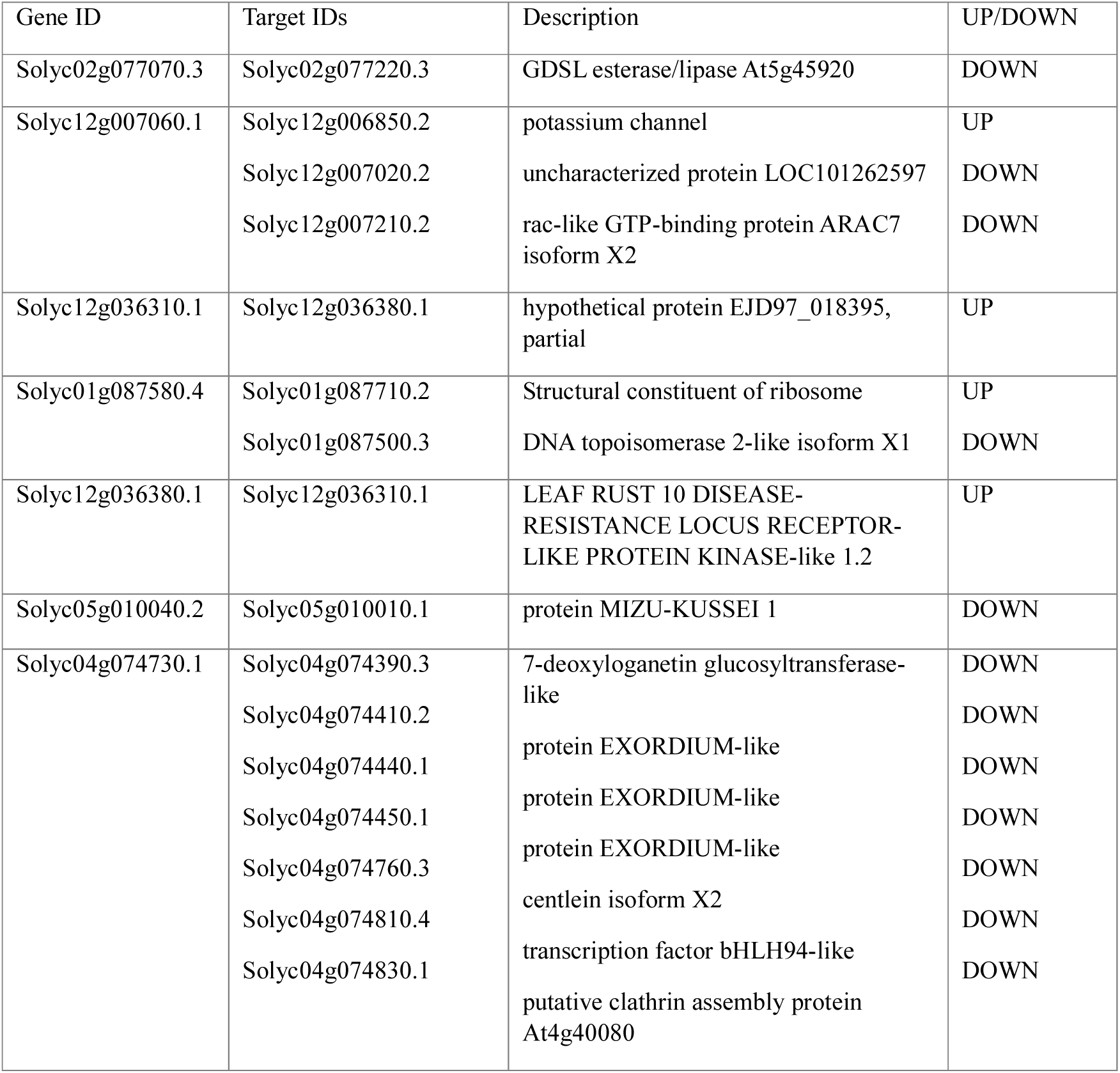
List of LncRNAs and their predicted *cis-*targets.

### LncRNAs and miRNAs

LncRNAs can act as the precursors of miRNAs and regulate the expression of various genes. In the present study, we identified two lncRNAs, one each in the leaf and root of tomato plants treated with low cadmium **(Table no.2)**. Solyc01g006780.4 was seen to be having a role in the expression of sly-MIR396a in *S. lycopersicum* and Solyc12g019150.1with ppt-MIR1063g (sly-MIR1063g) in *Physcomitrella* patens, in leaves and roots respectively. Based on our RNA-Seq data, these lncRNAs were differentially expressed in low cadmium-treated plants with a fold change of >1 and p-value <0.05. The relative fold change of lncRNAs was analyzed and confirmed using RT-PCR analysis. The transcript level of mi-RNAs was also confirmed using stem-loop RT-PCR **(Fig.7).** The putative targets of these mi-RNAs were predicted using the psRNATarget server and compared the targets to the differentially expressed genes from our RNA-Seq data. We found that 5 genes among the predicted targets by sly-MIR396a and one gene from the predicted targets by sly-MIR1063g were differentially expressed in RNA-Seq data **(Table no.3),** which was further confirmed by RT-PCR analysis. **(Supplementary Fig.3**).

**Fig. 7:**
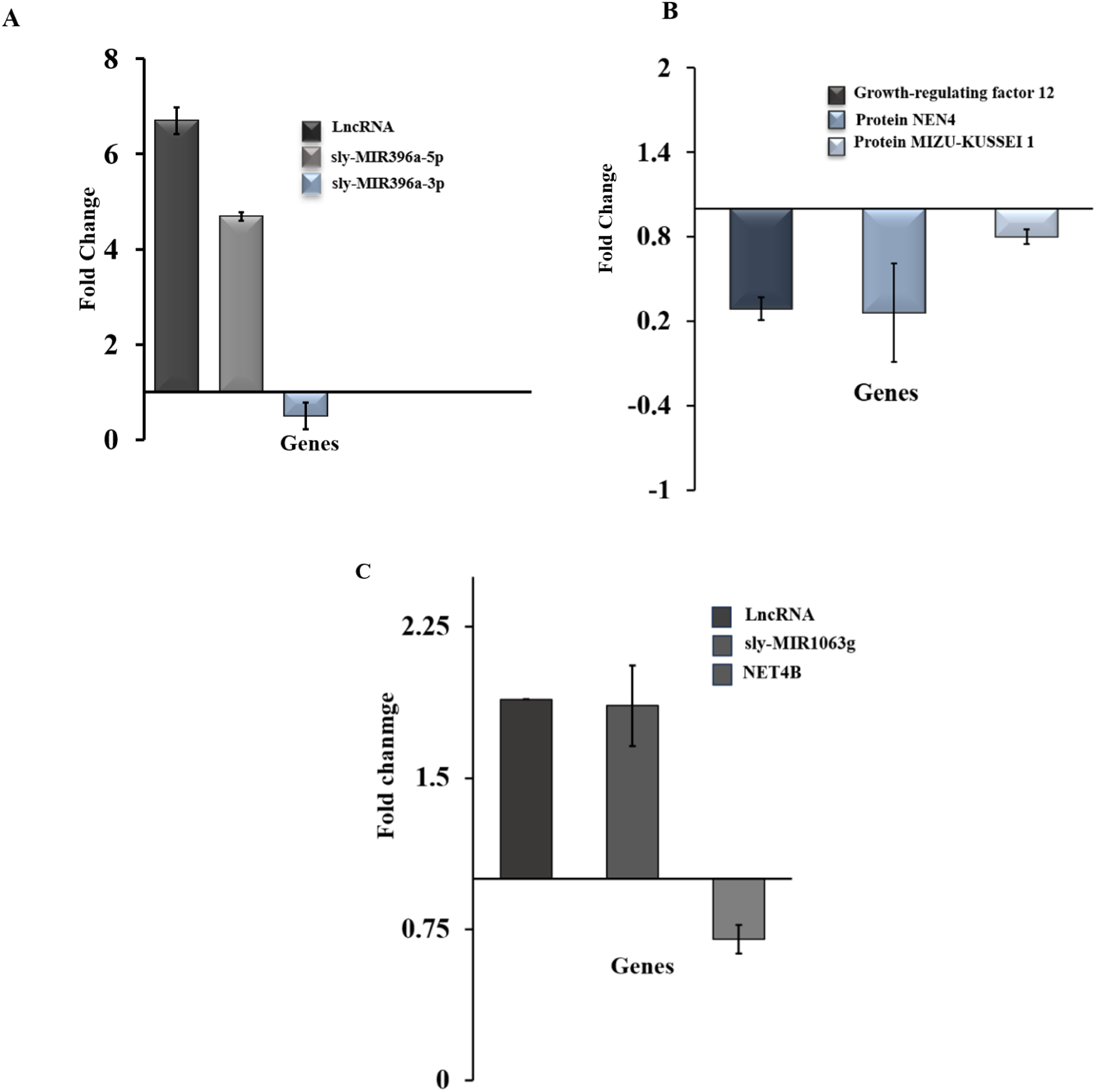
Real-time PCR analysis of LncRNAs, miRNAs, and miRNA targets in leaves and roots of tomato plants treated with 1µM Cd. A) Expression Fold change in the lncRNA, its miRNAs, and B) miRNA target genes in the leaves of tomato plants treated with 1µM Cd for five days. C) Expression Fold change of differentially expressed lncRNA, its miRNA, and miRNA target in the roots of tomato plants treated with 1µM Cd for five days. Bars in the graph represent the mean ± SD, calculated from duplicates.

### sly-MIR396a regulation of Growth-Regulating Factors (GRFs)

GRFs are known to be targets of sly-MIR396a, and to confirm their interaction, the relationship between GRFs and sly-MIR396a was investigated using the RNA-hybrid software. The sequences of all known GRFs in tomato and the mature sequence of sly-MIR396a were downloaded from Sol genomics and miRBase, respectively. They were uploaded to RNAhybrid software to check whether all GRFs are targets of sly-MIR396a **(Fig.8).** The results indicated that all the GRFs except Solyc08g068760.1.1, Solyc08g079800.2.1, and Solyc09009200.1.1 are potential targets of sly-MIR396a with a minimal free energy hybridization value which less than −33kcal/mol. The minimal free energy hybridization values for Solyc08g068760.1.1, Solyc08g079800.2.1, and Solyc09009200.1.1 were −18.0kcal/mol, −27.8kcal/mol and −31.3kcal/mol respectively which exceeded than the minimal values of most of the miRNA-target pairs and also the number of mismatches in this pairs were far more than the other pairs suggesting that theses GRFs may not be the potential targets of sly-MIR396a. The Solyc01g091540.1.1, which was upregulated in our RNA-Seq data of 1µM Cd-treated leaves, had a minimal free energy hybridization value of −33.9kcal/mol, which authenticates it as a possible target of sly-MIR396a.

**Fig. 8:**
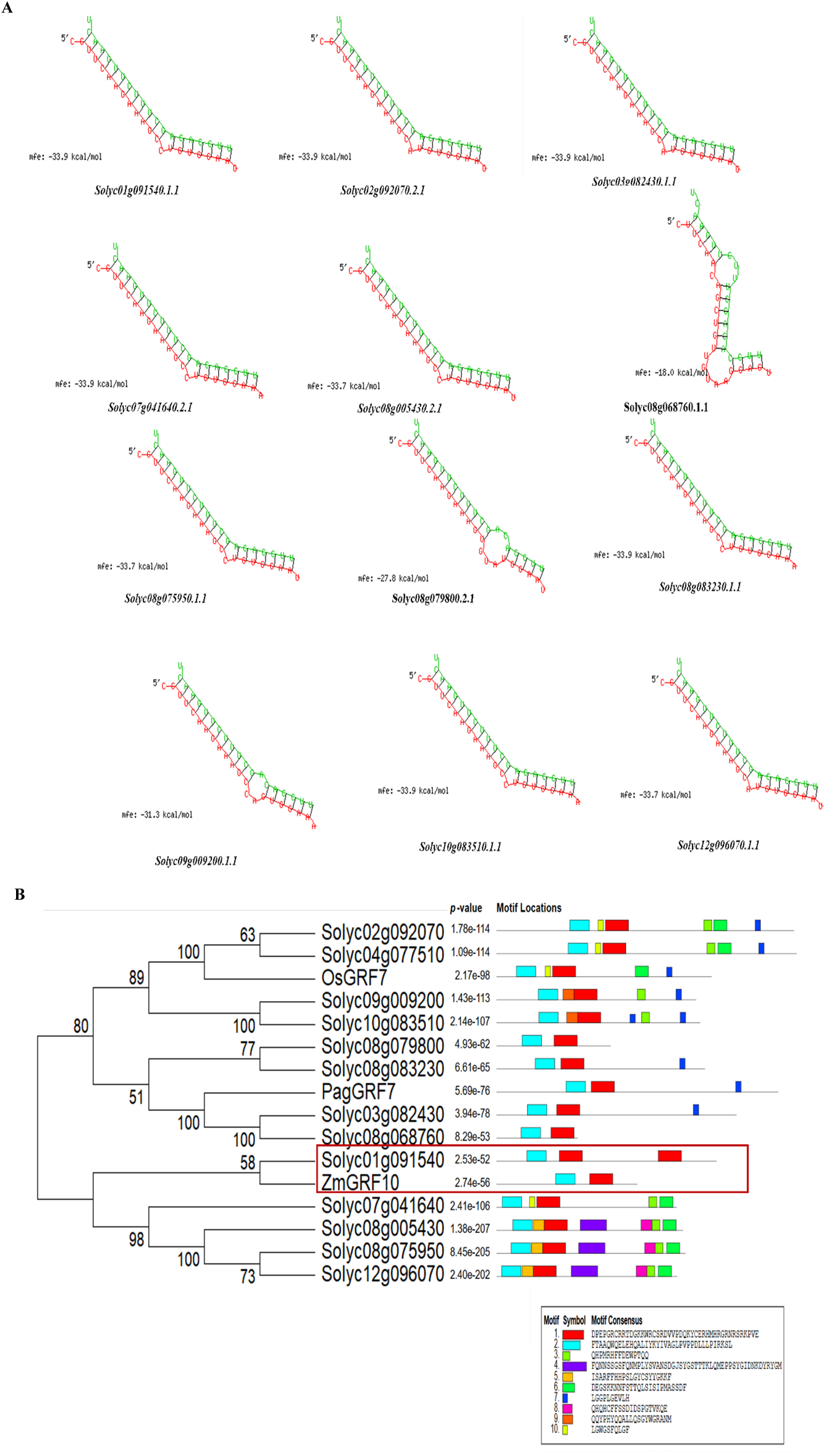
Interaction analysis between miRNA and targets. A) The minimum free energy hybridization between miR396a and GRFs in tomato calculated using RNAhybrid software. The red and blue colours represent the miRNA and targets, respectively. B) Functional and Phylogenetic similarities of GRF12 with other GRFs: Phylogenetic and gene structure analysis of GRFs. A neighbour-joining phylogenetic tree was constructed using MEGA12 after aligning full-length GRF sequences using the Clustal option. The conserved domains of GRF sequences were retrieved using the MEME Suite software. QLQ (Gln-Leu-Gln) and WRC (Trp-Arg-Cys) domains, FFD (Phe-Phe-Asp), TQL (Thr-Gln-Leu), and GPL (Gly-Pro-Leu) motifs are the common conserved domains and are represented in different coloured boxes

### Sly-MIR396a downregulates the expression of growth-regulating factors

GRFs interact with GIFs and regulate cell proliferation during leaf development. Most GRFs act as positive leaf cell size regulators, whereas some GRFs can reduce cell size. In this study, we found the GRF to be downregulated by the sly-MIR396a and the leaf size to be increased in the 1µM Cd condition. The phylogenetic analysis of GRFs in tomato, Arabidopsis, Maize, Rice, and Poplar was shown in **Fig.9**. The Solyc01g091540.1.1 (GRF12 according to Solgenomics) was seen to be closely related to ZmGRF10. Conserved domains and motifs were also investigated, and all the GRFs showed the characteristic QLQ and WRC domains in their N terminus. The leaf cell analysis of the control and hormetic conditions was performed to check the effect of downregulated GRF. The leaf area, cell area, and number of cells were measured using ImageJ software and DIC microscopy **(Fig.9).** The leaf area, cell area, and number of cells were greater in the hormetic condition compared to the control and stress conditions.

**Fig. 9:**
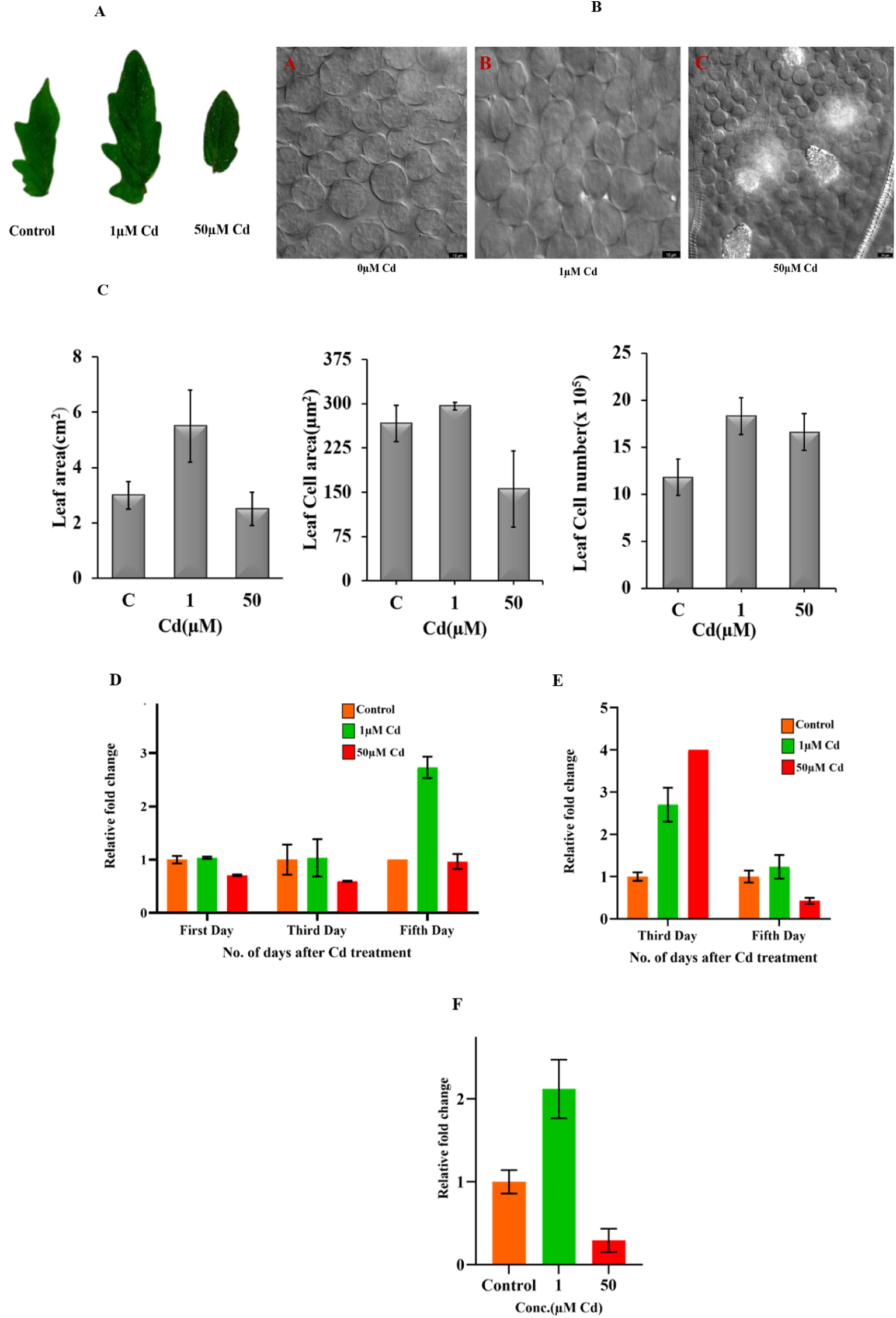
Microscopic images of leaves by DIC. B) Palisade cells in control, 1µM, and 50µm cadmium-treated plants. Bar = 10µm. After treatment, the third completely open leaves were collected and fixed in FAA solution, after which they were treated with clearing solution overnight and observed under a differential interference contrast (DIC) microscope. C)The leaf area, Leaf cell area, and Leaf cell number were calculated using Image J software. (D-F) Gene expression analysis of cell cycle and cell expansion genes. RT-PCR analysis of D) Cyclin P3-1-like, E) Wall-associated receptor kinase 2-like, and F) Endoglucanase 21-like genes. Expression Fold change of genes in the leaves of tomato plants treated with 1µM Cd for five days. Bars in the graph represent the mean ± SD, calculated from duplicates. Ubiquitin was used as the internal control.

### Sly-MIR1063g activates root growth by regulating NET4 B expression

The interaction between the **S**ly-MIR1063g and NET4B (NET4B is a protein belonging to the NETWORKED (NET) family, known for its role in actin-membrane) interactions was analysed using RNAhybrid tool, and the minimal hybridisation value was obtained as −24.7kcal/mol (**Fig.10).** This, along with the mismatches, indicates that miRNA and NET4B have a moderate level of interaction but still is considered as a possible target of the miRNA. The root tips were visualised under the DIC microscope, and the number of cells and cell size were observed to be higher in roots treated with 1µM Cd condition, where the NET4B was seen to be downregulated by **S**ly-MIR1063g. DIC

**Fig. 10:**
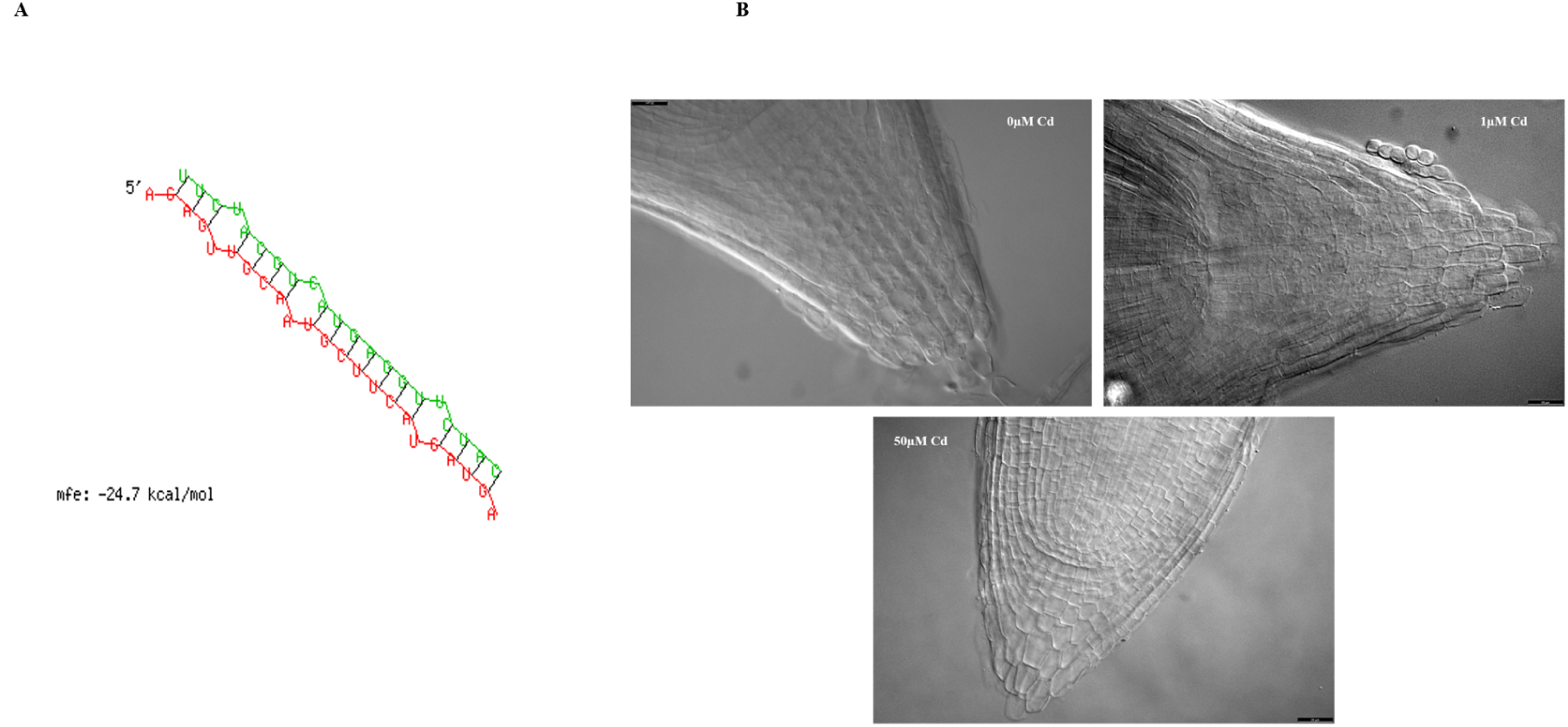
Interaction analysis between miRNA and NET4B. A) The minimum free energy hybridization between miR396a and GRFs in tomato was calculated using RNAhybrid software. The red and blue colours represent the miRNA and targets, respectively. Microscopic images of the root by DIC. B) Root cells in control, 1µM, and 50µm cadmium-treated plants. Bar = 10µm. After treatment, the root tips were collected and fixed in FAA solution, after which they were treated with clearing solution overnight and observed under a differential interference contrast (DIC) microscope.

## Discussion

Hormesis is a dose-dependent response exhibited by plants where the lower dose has a stimulatory and the higher dose has an inhibitory effect on the growth and development of plants (Yang et al., 2019). The detrimental chemical agent or environmental factor at a higher concentration induces an adaptive beneficial effect to the plant. Although all toxic substances do not initiate a biphasic response, most of them exhibit this feature, which is a clear indication of hormesis. In most of the studies, it was observed that the hormetic phenomenon does help organisms tolerate more than one type of stress (Mattson., 2007). In hormetic responses, plant growth is regulated by the production of auxins, anatomical variations, etc. and also through the production of anti-oxidant enzymes. Different hormetic features inducing several independent cellular functions, along with multiple signals and receptors produce an integrated cellular response leading to increased plant growth and development. Hormesis has been adopted and applied in medical, biological, and agricultural areas (Jalal., 2021). Hormesis in animals usually involves activating the production of growth factors, transcription factors related to stress-related proteins like HSPs, anti-oxidant pathways, ion channels, different kinases, etc. (Vargas-Hernandez et al.,2017). In case of plants, the exact mechanism behind hormesis is still unknown. However, a lot of possible explanations have been proposed which include 1) hormetic considered as the growth stimulant, 2) an indirect mechanism involving over compensation response activating plant defence 3) metal hormesis inducing the plant stress tolerance mechanisms, and 4) generation of minimal ROS leading to anti-oxidant defense, signaling and ultimately growth stimulation which is considered being the most plausible mechanism (Vargas-Hernandez et al.,2017).

Cadmium, a highly toxic heavy metal readily absorbed by plants has always gained much attention due to its long biological half-life and toxicity in relatively small quantities. In plants, cadmium stress causes reduced photosynthesis and yield, stunted growth, DNA damage, membrane disintegration, etc. However, the stimulatory effect/hormetic responses of cadmium on plants in sublethal doses have recently attracted scientists’ interest. In the present study, we could observe that the growth parameters like root length, shoot length, fresh weight, and dry weight of tomato seedlings grown under low doses of cadmium were significantly increased compared to control and higher doses of cadmium. These findings generated a ‘U’ shaped or inverted ‘U’ shaped graph confirming the hormetic responses of tomato seedlings. Similar findings were reported by Liu et al., 2015 and Liu et al., 2023, in *Lonicera japonica* and in Barley by Aery and Rana (2003). Among the different cadmium concentrations, 1µM Cd showed maximum growth, and 50µM Cd showed severe growth inhibition.

Apart from physiological criteria biochemical parameters like proline content, MDA content, RLWC, and anti-oxidant enzyme activities were also investigated under different doses. Proline is a common amino acid encountered in plants and an osmolyte that accumulates in various abiotic stresses (Szabados and Savouré, 2010; Slama et al., 2015). Proline also helps plants scavenge ROS, stabilize proteins and membranes, and regulate cellular redox potential under various stress conditions (Ashraf and Foolad,2016). Accumulation of proline; a common physiological response in plants exposed to heavy metal stress can be due to *de novo* synthesis, lower utilization, protein degradation, or reduced degradation (Kaur and Asthir, 2015). We observed no significant change in proline content in the leaves of plants treated with lower cadmium concentrations, whereas in 50 µM Cd treated plants a significant amount of proline was accumulated compared to control. However, in the case of roots, there was a significant decrease of proline in 1 µM Cd-treated plants and a profound increase in 50 µM treated plants. The reduced amount of proline in 1µM Cd either indicates that cadmium exposure at this dose does not induce stress in plants to accumulate proline or the proline produced might be utilized for other cellular processes like a heavy metal chelator, signaling molecule, or a gene regulator which requires further clarification. The MDA or malondialdehyde another stress marker is one of the final products of lipid peroxidation of polyunsaturated fatty acids. It is a highly reliable marker to determine the degree of damage incurred by stress to the plant (Morales and Munné-Bosch, 2019). Lipid peroxidation can damage the proteins of cell membranes thereby severely compromising the membrane integrity and even leading to the death of plant cells in extreme cases (Zhang et al, 2021). Thus, the higher the MDA content higher the damage from biotic or abiotic stress (Alché, 2019). The MDA in 1 µM leaves was not significantly different from the control and the MDA content was significantly higher than the control in 50 µM. The values suggest that 1µM Cd exposure does not generate oxidative damage or stress in plants. These findings concur with the previous studies done on peppermint and tomato seedlings where the MDA content was reduced when treated with low doses of cadmium and glyphosate respectively (Wang et al. 2023; Wang et al. 2024). Interestingly, our results showed no significant change in the MDA content in the roots of Cd-treated plants despite roots being the primary route for the entry of Cd. RLWC is a measure of plant water status in terms of the physiological consequence of cellular water deficit. Water potential as an estimate of the energy status of plant water is useful in dealing with water transport in the soil-plant-atmosphere. The RLWC percentage we obtained in our experiments was similar to the observations made by Liu et al. 2015, in *Lonicera japonica* plants, where the RLWC was significantly increased in lower cadmium concentrations and was decreased in higher cadmium exposure. This indicates that Cd is affecting the water balance of the plant in a way that the low doses exert a positive influence.

Thus, in light of all these observations, we concluded that low cadmium has beneficial effects on the growth and development of plants, and 1µM Cd can be considered the optimum dose for inducing cadmium hormesis in tomato plants. In the background of these promising results, we decided to check whether the hormetic effects persist even after the removal of cadmium from the medium. To achieve this, after five days of treatment with 1µM Cd, we removed and replaced the media of one set of plants with normal half-strength Hoagland’s media and checked the growth for one month. The growth parameters like length and biomass, were measured every three days for one month and observed that the plants with devoid of cadmium still showed a significant increase in length and biomass compared to the control. Another promising observation was that the growth of plants with cadmium removed was comparable with the plants subjected to continuous exposure of 1µM Cd, demonstrated the beneficial effects of hormesis, and it can persist even with acute exposure to the hormetic agent, which can indeed revolutionize the practical utility of this phenomenon.

To get a more elaborate idea of this particular mechanism, transcriptomic profiling of the plants has been performed with 1µM Cd as the hormetic condition and 50µM Cd as the stress condition against control (0µM Cd). The number of differentially expressed genes (DEGs) found in leaves was more than double the number of genes in roots. Also, the number of DEGs found in 50 µM Cd is far higher than the genes in 1µM Cd condition. The upregulated differentially expressed genes unique to 1µM Cd leaves are mainly involved in oxidoreductase activity, transport, and photosynthesis. In the case of roots, they are mostly secondary metabolite synthetic genes, detoxification proteins, and transporters. However, in 50 µM Cd condition, upregulated genes in leaves include transcription factors, DNA repair genes, jasmonic acid-related genes, and respiratory burst oxidase homolog protein. In roots, most of the upregulated genes belong to cell wall synthesis as expected in cadmium stress conditions.

In an attempt to delineate the molecular mechanism of hormesis, we explored the DEGs in 1µM Cd condition. Apart from the genes mentioned, several long non-coding genes were seen to be differentially expressed in the leaves and roots of 1µM Cd condition. They are transcripts with more than 200 nucleotides long and do not code for any proteins. They interact with mRNA, DNA, miRNA, and proteins and regulate gene expression at translational, transcriptional, or post-translational levels (Zhang et al. 2019). After analysing the coding potential of the lncRNA candidates using CDC 2.0 and PNRD software, those having a potential <1 or negative value were selected for further studies (Supplementary Table.1). The *cis-*targets of selected lncRNAs were compared with RNA-Seq data to determined that the functional aspects of lncRNAs. Among the targets, observed that potassium transporter genes, gibberellin and ethylene synthetic genes, and ALP-1 (Alkaline phosphatase 1) genes were upregulated, and DNA topoisomerase-like protein, EXORDIUM-like protein, aquaporin, and peroxidase were downregulated and revealed that the low Cd exposure leads to lncRNAs involved in transporter activity, hormone synthesis, chromatin remodelling, DNA replication, etc.

The lncRNAs, apart from directly interacting with proteins and DNA, they can also act as precursors of miRNAs. miRNAs are a class of small RNAs that share high complementarity with the target mRNAs and regulate their expression. They play an important role in plant development by responding to abiotic/biotic cues (Miller, 2020). There have been a lot of reports on the role of lncRNAs and as miRNA precursors in conferring tolerance to cadmium stress in different plants (Feng et al.2016; Wen et al. 2020; Li et al.2023; Qiu et al. 2024; Li et al.2024). However, lncRNAs as miRNA precursors in cadmium hormesis is unexplored. Two lncRNAs were identified as miRNA precursors and their 1µM Cd condition in each leaf and root. The miRNAs were predicted using miRBase software and their targets by psRNATarget software and were compared with RNA-Seq data, and found that in the leaf, Solyc01g006780.4 was the predicted precursor of sly-MIR396a in *S. lycopersicum.* The sly-MIR396a is a conserved miRNA in the plant kingdom, which helps in the growth and development of plants and also mitigates abiotic stresses through the regulation of the expression pattern of different targets (Yuan et al. 2020). The predominant target of this miRNA is a group of plant-specific transcription factors called Growth Regulation Factors (GRFs) (Mecchia et al. 2012). The Sly-MIR396a is upregulated along with the precursor Solyc01g006780.4 and confirmation by RT-PCR suggests the role of Sly-MIR396a in increased growth during hormesis.

According to the predictions by psRNATarget (Table 2), all the targets obtained were down-regulated in RNA-Seq. Among the targets is Solyc01g091540 (GRF12 according to the Sol genomics database), a growth-regulating factor that seems to have some role in cadmium hormesis. Wu et al (2022) reported that GRFs are important transcription factors specific to plants, regulating root and stem growth, leaf size, longevity, and flowering and have a highly conserved QLQ and WRC domain in the N terminus and a highly variable C terminus. The QLQ domain interacts with GIFs (GRF Interacting Factors) and WRC is a DNA interacting domain (<u>Kim and Kende, 2004</u>; <u>Kim *et al*., 2012</u>). The C terminus is involved in transactivation and those GRFs with truncated C terminus do not exhibit the transactivation activity (<u>Choi *et al*.,</u> <u>2004</u>; <u>Liu *et al*., 2014</u>). Almost 13GRF genes have been characterized so far in tomato plants (Khatun et al. 2017). The differential expression of various GRFs in different stresses like heat, cold, UV-radiation, etc., indicates the function of these transcription factors in stress management in addition to growth regulation (Khatun et al. 2017). GRFs are predominantly found in young active regions of plants and are involved in cell proliferation. and reported that the elevated expression of GRFs increases cell size, leading to leaf expansion (Kim et al., 2003). However, studies revealed that even single GRF loss-of-function mutants have a minimal impact on leaf growth (Lazzara et al. 2024). It is reported that *ZmGRF10* in maize causes a reduction in the leaf size and plant height (Wu et al.2014) and AtGRF9 negatively regulates the cell proliferation and leaf expansion in Arabidopsis plants (Omidbakhshfard, et al.2018). Thus, some GRF factors might be acting as dominant negative variants and this allows us to hypothesize that lower expression of GRF12 produces plants with expanded leaves and increased height (Lazzara et al. 2024). Recent studies on GRF showed that they can coordinate plant growth with signaling and stress response. Under normal conditions, plants envisage various strategies to repress the expression of genes related to stress response. The AtGRF7 gene for example suppresses the stress-responsive genes under normal conditions. On the contrary, the genome profiling of the *atgrf7-1* T-DNA insertion mutant has disclosed several upregulated genes expressed in response to stress (Omidbakhshfard., et al.2018).

**Table no.2:**
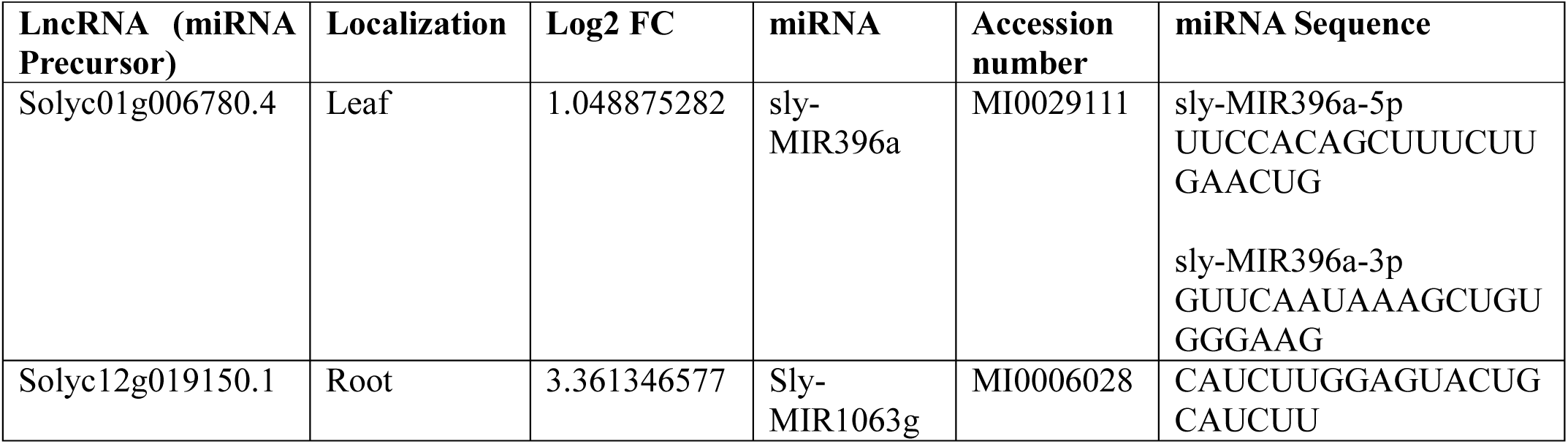
List of LncRNAs as precursor miRNAs along with their predicted miRNAs.

**Table no.3:**
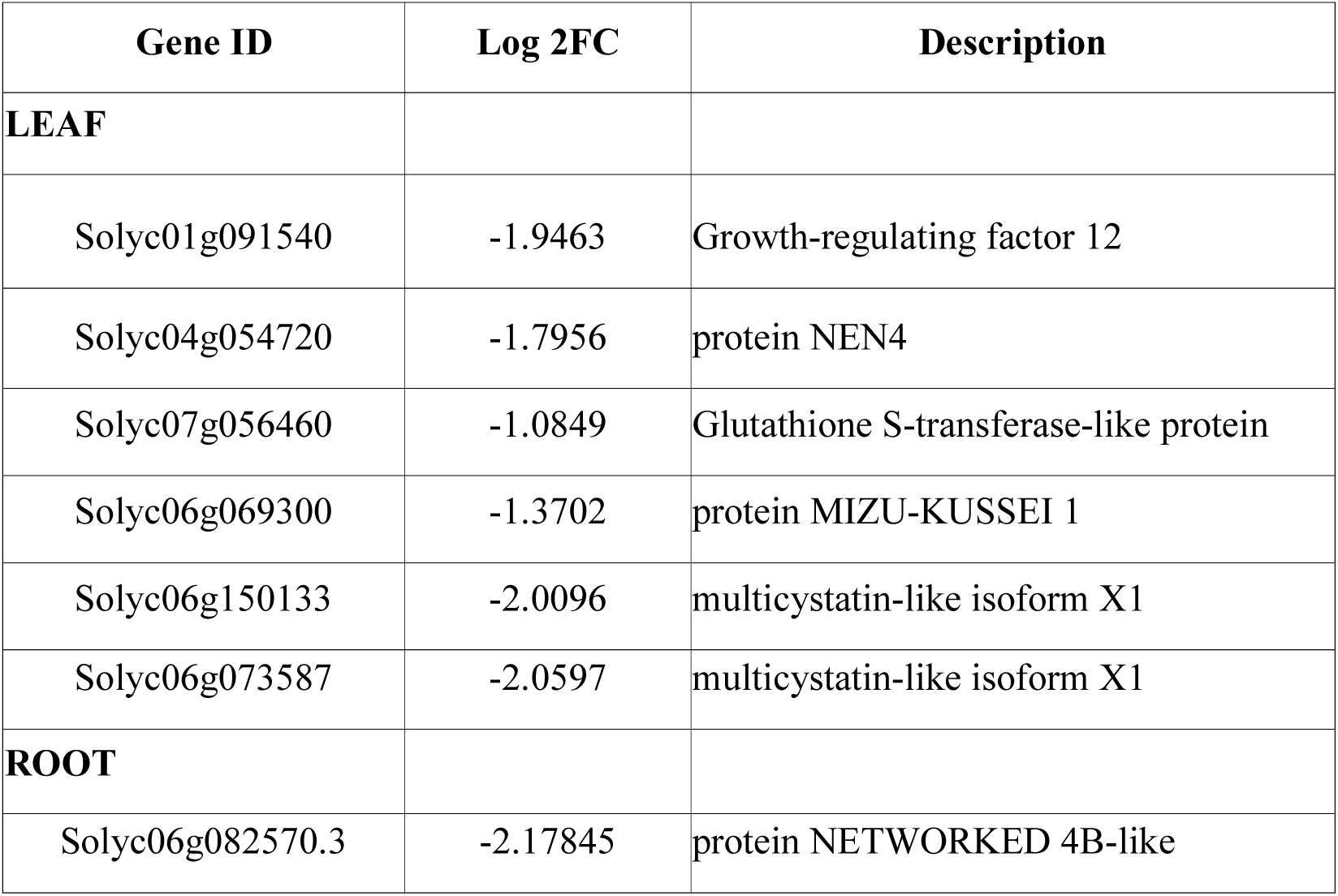
List of miRNAs and their predicted targets using psRNATarget software.

Comparison of GRFs we could infer that GRF12 of tomato closely related to ZmGRF10, that negatively regulates leaf size in maize plants where it lacks the truncated trans-activation domain, which is characteristic of ZmGRF10, indicated that the both these genes might behave differently (Wu et al.2014). According to Wu et al.(2014), the ZmGRF10 controls leaf size by regulating the expression of cell cycle genes and further reported that overexpression of ZmGRF10 in maize has led to the downregulation of cyclin genes, especially cyclin B2:1, resulting in the reduction of leaf size. In contrast, we have seen that cyclin B2 is downregulated in hormetic conditions, which suggests the involvement of other genes responsible for cell proliferation in hormesis. The Cyclin-P3-1-like (Solyc01g098930.3) is upregulated in leaves treated with 1µM Cd, which belongs to the P-type cyclin family, specifically expressed in root meristem, lateral caps, and responsible for the cell division in meristem (Chen et al., 2020). They activate the cyclin-dependent kinase B2-1 (CDKB2;1) and promote cell division during the G2-M phase of cell division, similar to Cyclin B (Lee et al., 2003; Ito, 2000). Hence, we hypothesise that the upregulation of Cyclin-P3-1-like (Solyc01g098930.3) in the leaves has lead to the activation of cell proliferation in tomato plants treated with 1µM Cd. In microscopic analysis, apart from cell number, cell size was also larger in the leaves of the hormetic condition compared to the control and stress conditions. The expression of cell expansion genes like endoglucanase 21-like (Solyc04g064900.1), and wall-associated receptor kinase 2-like (Solyc09g014720.3) in the leaves of 1µM Cd plants aligns with this observation (Kohorn and Kohorn, 2012; Wagner and Kohorn, 2001; Glass et al., 2015).

The expression analysis of Cyclin-P3-1-like and wall-associated receptor kinase 2-like genes was analysed on the first, third, and fifth day of cadmium treatment **(Fig.9).** As the leaf development is a complex process where cell expansion and proliferation occur in a well-coordinated manner during organ development, cell proliferation occurs first, followed by cell expansion and differentiation. Once the cell proliferation ceases, the cell starts to expand in size (Ezaki et al., 2024). Following our observation, in the hormetic condition, the expression of the cell proliferation gene ‘Cyclin-P3-1-like’ was not significantly different from control on the first and third day of cadmium treatment on the fifth day, there was significant up-regulation in that particular gene compared to the control. Also, the cell expansion-related gene ‘wall-associated receptor kinase 2-like’ was significantly upregulated on the third and fifth day of treatment in the hormetic condition compared to the control. Hence, from the above analysis, we were unable to screen out the exact dynamics of leaf growth in hormesis; however, we were able to identify that both ‘Cyclin-P3-1-like’ and ‘wall-associated receptor kinase 2-like’ play a key role in leaf growth during cadmium hormesis, which requires further investigation. Thus, we can infer that the GRF12 probably represses the expression of stress-responsive genes, is downregulated by Cd exposure, and the stress-responsive genes are expressed within the threshold level. This sub-minimal level of expression of stress-responsive genes gave rise to hormetic stimulation, which in turn is based on the moderate activation of random defense mechanisms by plants that protect them from severe environmental stress with the involvement of chemical compounds, stress hormones, and the anti-oxidant defense system, as well as the synthesis of stress proteins (Agathokleous et al.2020).

The lncRNA (Solyc12g019150.1) upregulated in roots showed similarity with the stem-loop sequence of another miRNA known as sly-MIR1063g. This miRNA, originally identified in *Physcomitrella patens,* was predicted by blasting the lncRNA sequence against the miRBase database and further validated by RT-PCR analysis (Axtel et al., 2007). After careful prediction and comparison of the targets with the RNA-Seq data, we found that it suppresses the expression of the NET4 B-like protein in the roots of tomato plants. The Networked family of proteins is an actin-binding plant-specific protein family characterized by a NET actin-binding domain (NAB). Phylogenetically, protein is classified into four families: 1-4 (Hawkins *et al*.,2014). These proteins link the actin cytoskeletons to various membrane compartments. Studies have already shown the association of NET4A/4B proteins with actin filaments and the tonoplast membrane of the nucleus, and also established their potential to remodel the actin cytoskeletal organization and influence vacuolar morphology (Hawkins et al., 2023). The deregulation and overexpression of NET4A protein expression lead to the increased and decreased vacuolar occupancy observed in plant cells. In other words, the increased abundance of NET4A proteins can induce more compact vacuoles, which explains the reduced cellular and organ growth observed in plant cells.

NET4B protein was one of the targets predicted by the psRNATarget server for the sly-MIR1063g miRNA expressed in root cells of low Cd-treated plants. The gene was downregulated in roots, and the precursor lncRNA responsible for this miRNA was upregulated. NET 4B is the closest homologue of NET4A, and both co-localize to actin filaments and tonoplast (Hawkins et al., 2023). After correlating the above results with the previous experiments and observations, we conclude that the lower abundance of NET4B proteins can lead to less compact vacuoles and result in increased growth of plant cells, which can be in consonance with the improved root length exhibited by the plants treated with 1µM Cd. In conclusion, as shown in Figure 11, the miRNAs, sly-MIR396a, and sly-MIR1063g downregulate GRF12 and NET4B proteins, respectively leading to the increased growth observed in plants treated with 1µM Cadmium.

**Fig. 11.**
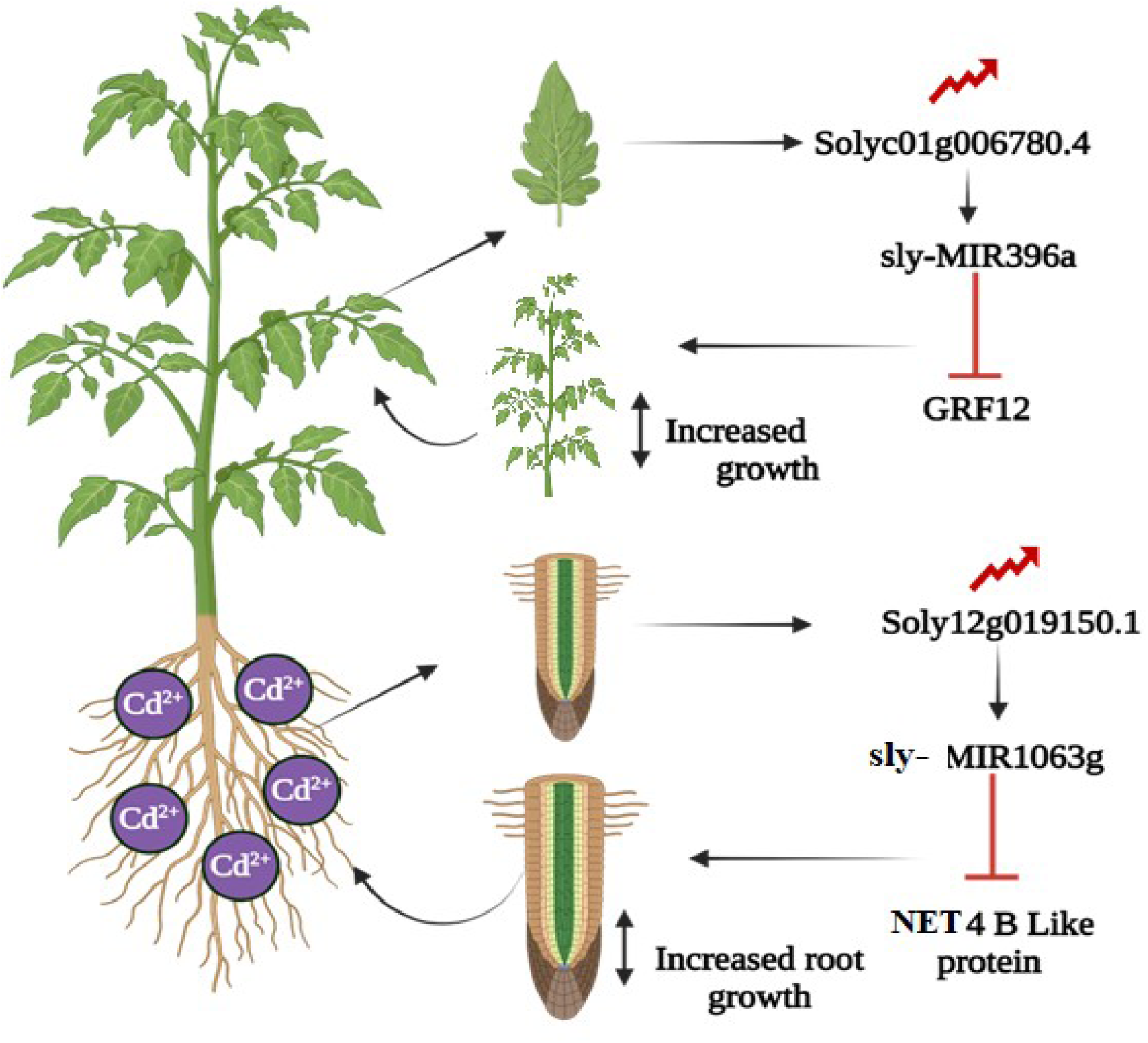

## Author contributions

RR: Performed the experiments and wrote the manuscript; MY: Performed experiments and helped in analysing results. SRK: Conceptualized, designed the experiments, interpreted the data, supervised the entire work, and edited the manuscript.

## Conflict of Interest

None

## Acknowledgement

We acknowledge the financial support of Science and Engineering Research Board (SERB) Govt. of India (EEQ/2020/000395) and acknowledged DST-FIST, UGCSAP, DBT-BUILDER infrastructural facilities of the department. RR and MY thankful to CSIR and UGC Govt. of India, for Research fellowship. We sincerely thank Dr.Pinninti Malathi, and Prof. Yelam Sreenivasulu for helping microscopic images

## Supplementary Tables

**Supplementary Table No. 1:**
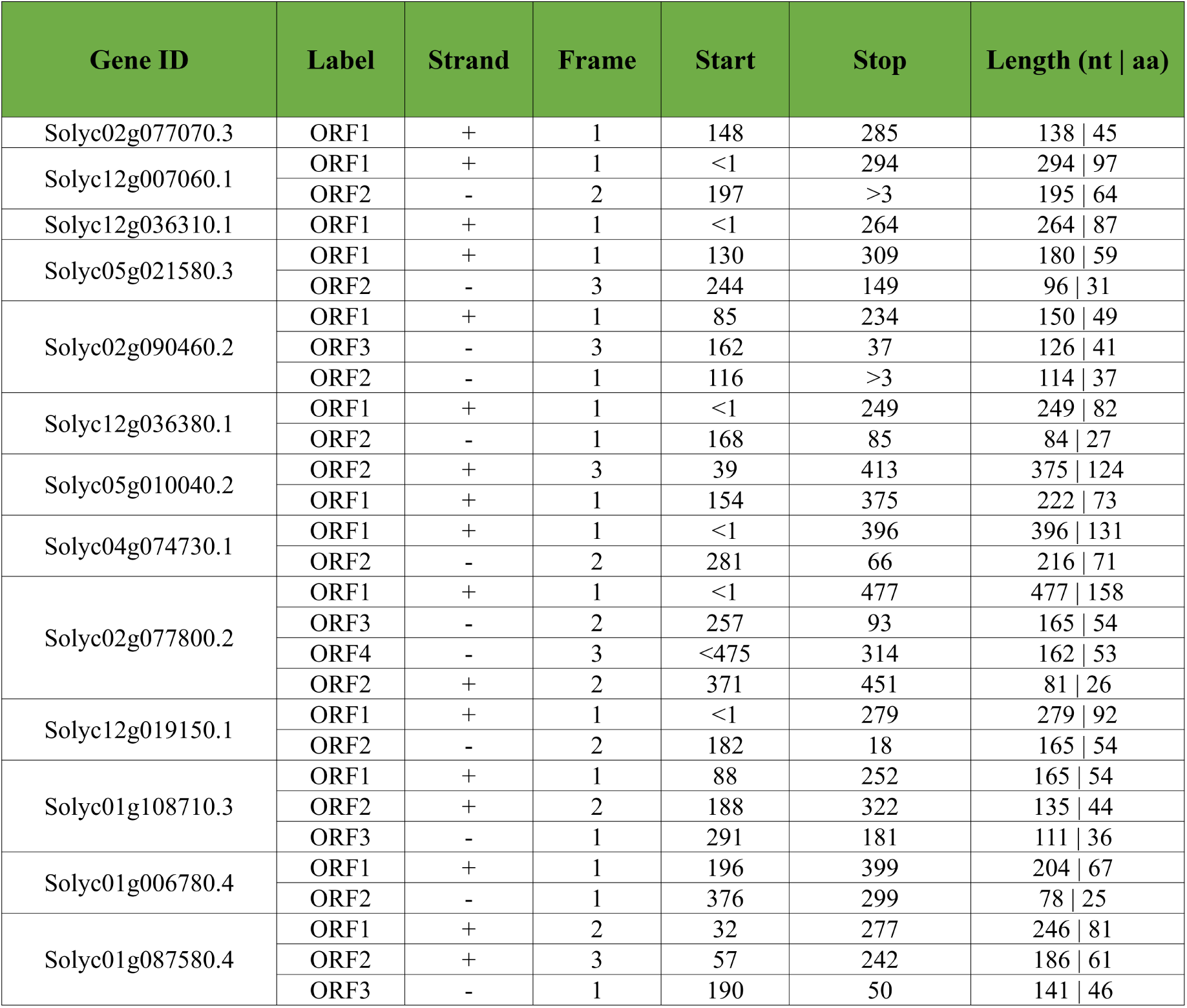
List of potential LncRNAs and their ORFs predicted using NCBI ORF Finder.

**Supplementary Table No. 2:**
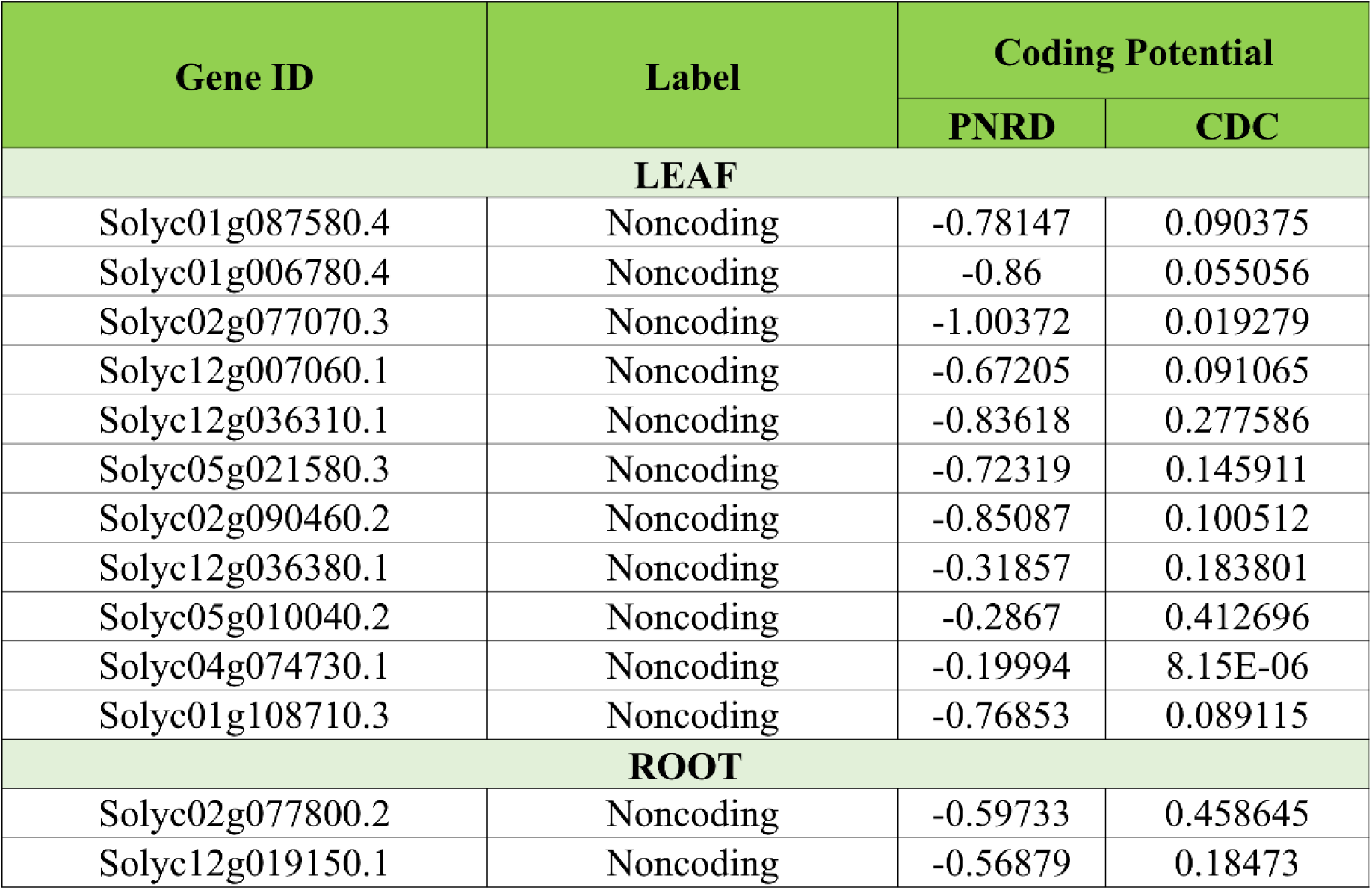
List of potential LncRNAs along with their coding potential.

**Supplementary Table No. 3:**
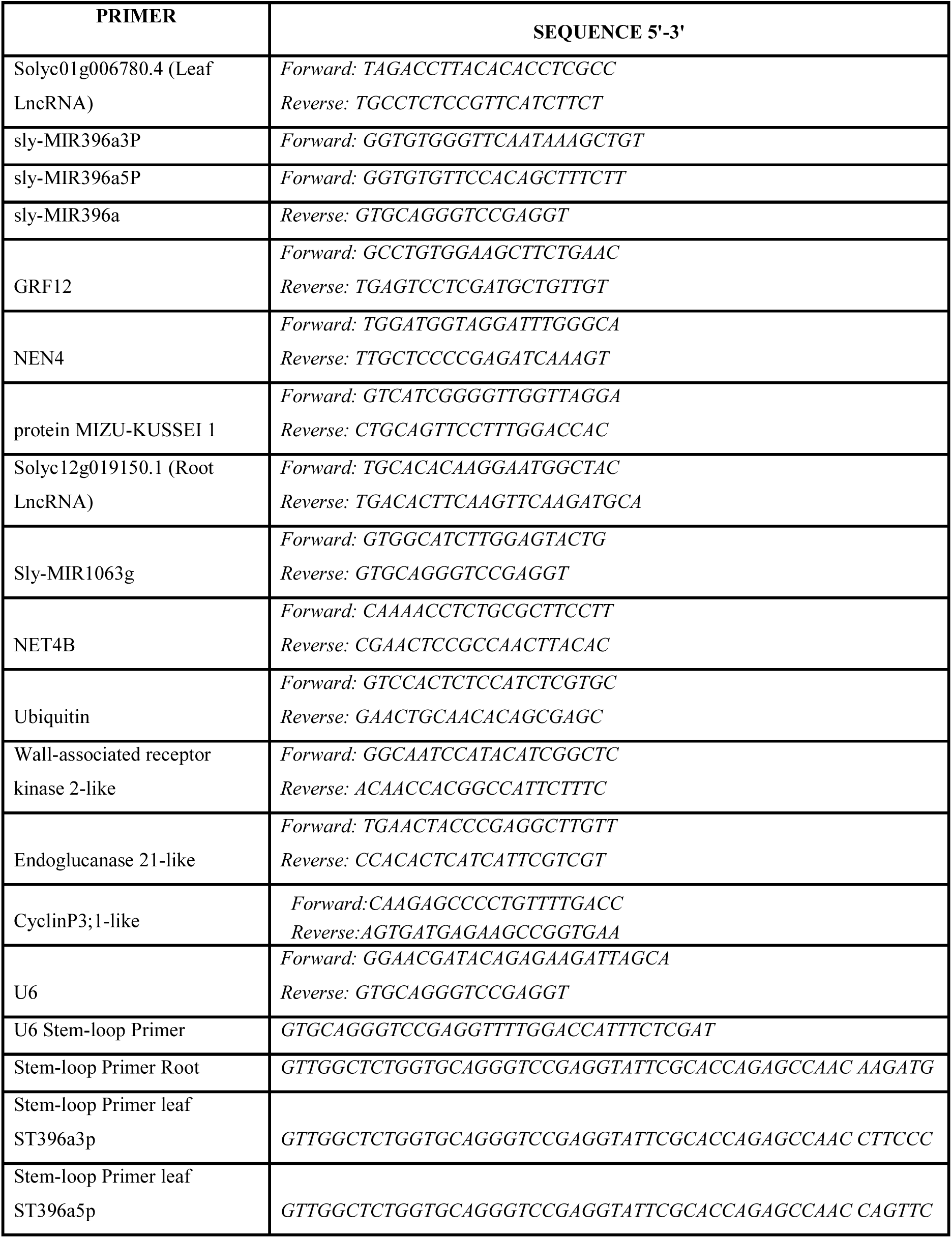
List of Primers used.

## Supplementary Figures

**Supplementary Fig. 1:**
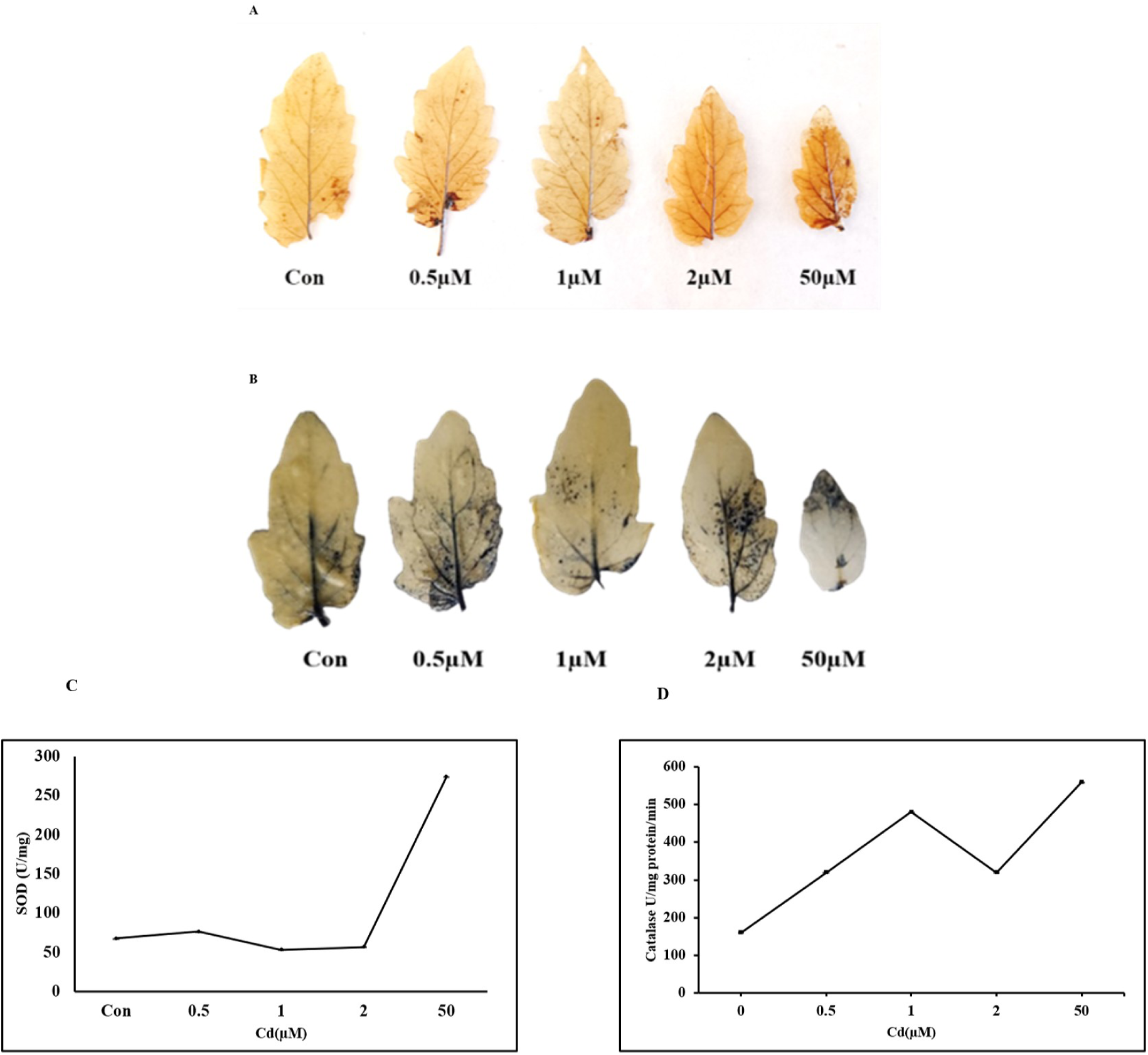
Histochemical analysis for measuring ROS in tomato leaves treated with cadmium for five days. A) DAB staining of tomato leaves treated with different cadmium concentrations. The dark brown colour in the leaves is produced by the oxidation of DAB in the presence of peroxidase and hydrogen peroxide. B) NBT staining of tomato leaves treated with different concentrations of cadmium. The blue colour represents formazan, produced from the reduction of NBT by free oxygen radicals. C) SOD and D) CAT enzyme activity in leaves of 20 days old tomato seedlings treated with cadmium for five days.

**Supplementary Fig.2:**
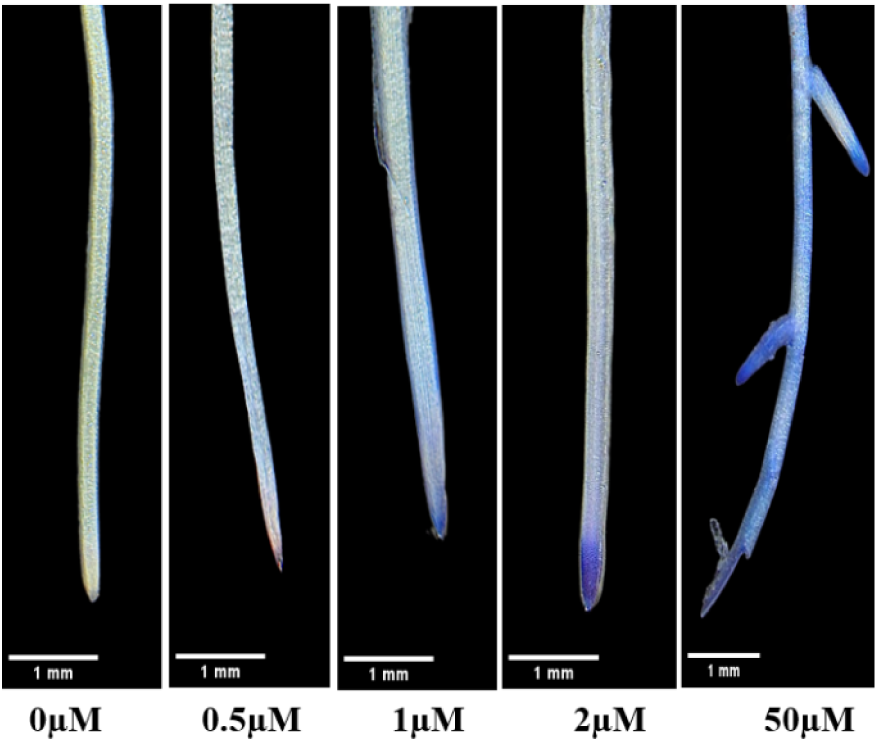
Evans blue staining of roots of tomato plants treated with different cadmium concentrations. The semi-permeable nature of cell membranes excludes the dye, whereas damaged cells or membranes with compromised integrity take up the dye and are stained blue.

**Supplementary Fig.3:**
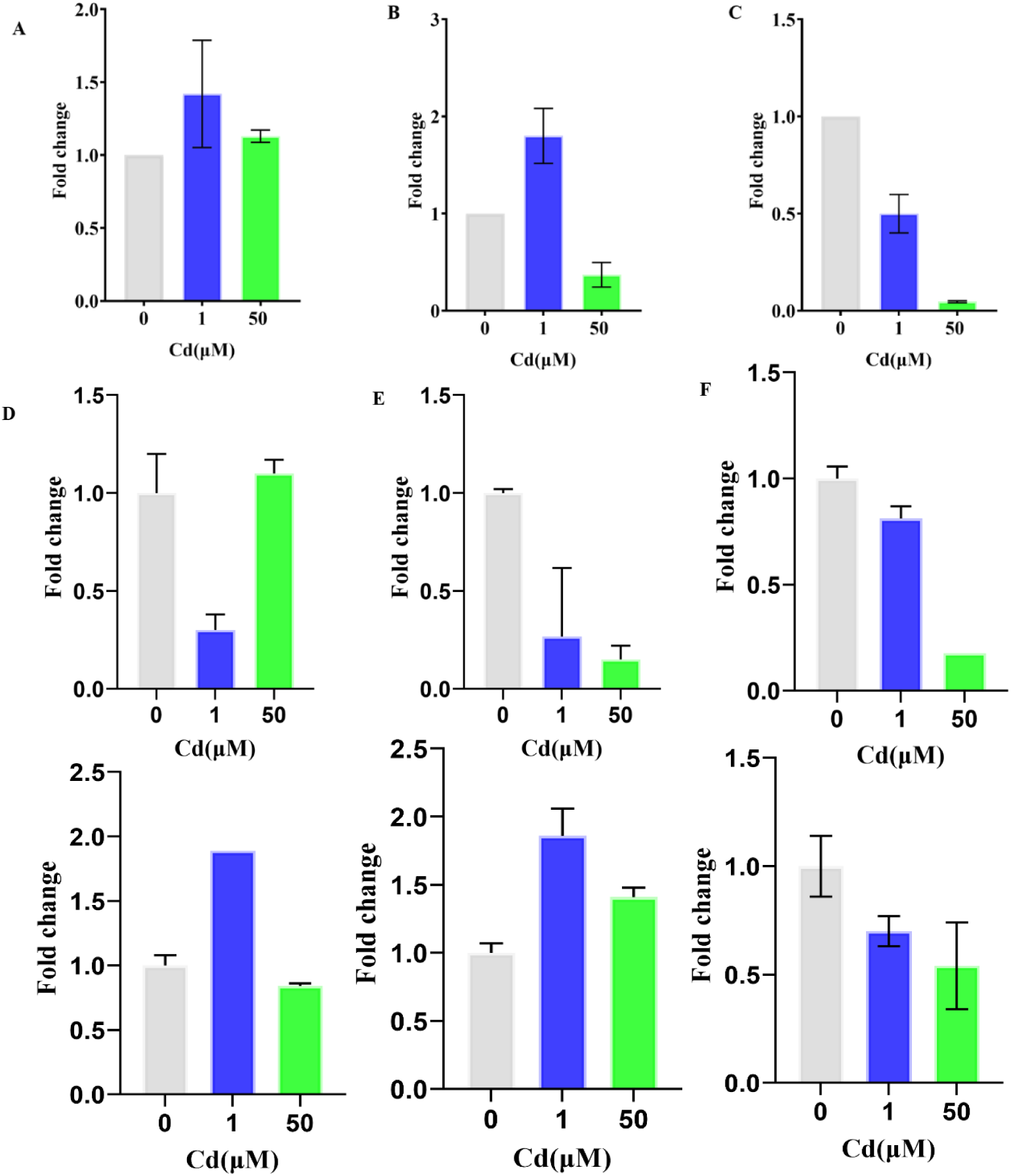
Real-time PCR analysis of A) LncRNAs, B) sly-miR396a-5p, C) sly-miR396a-3p, D) GRF12, E) protein MIZU-KUSSEI 1 F) NEN4 Protein in leaves and G) LncRNA, H) sly-MIR1063g, and I) NETWORKED4B Like protein in roots of tomato plants treated with 1µM Cd solution for five days. Actin and ubiquitin were used as the internal controls. Bars in the graph show mean ± SD calculated from duplicates.

**Supplementary Fig.4:**
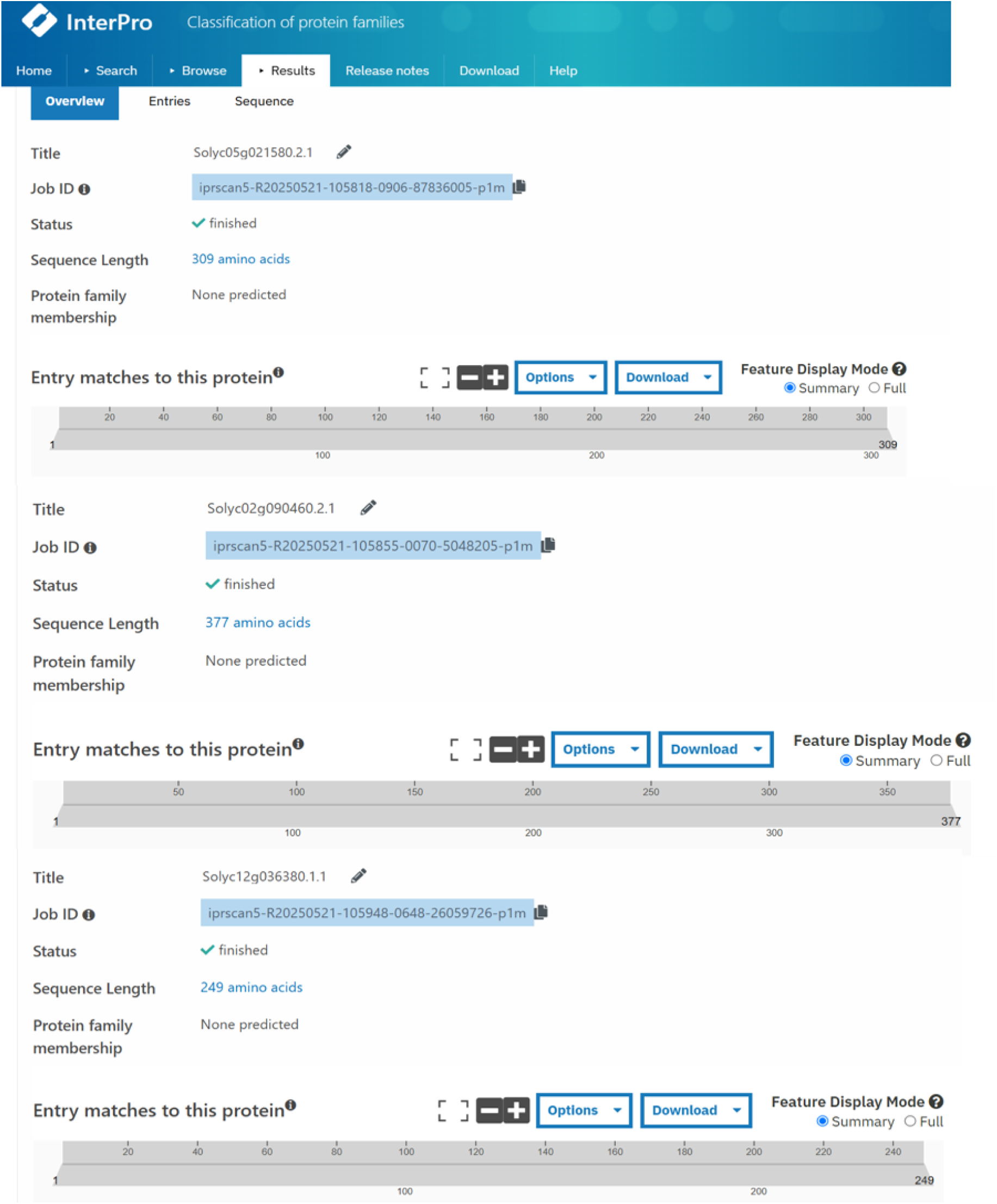
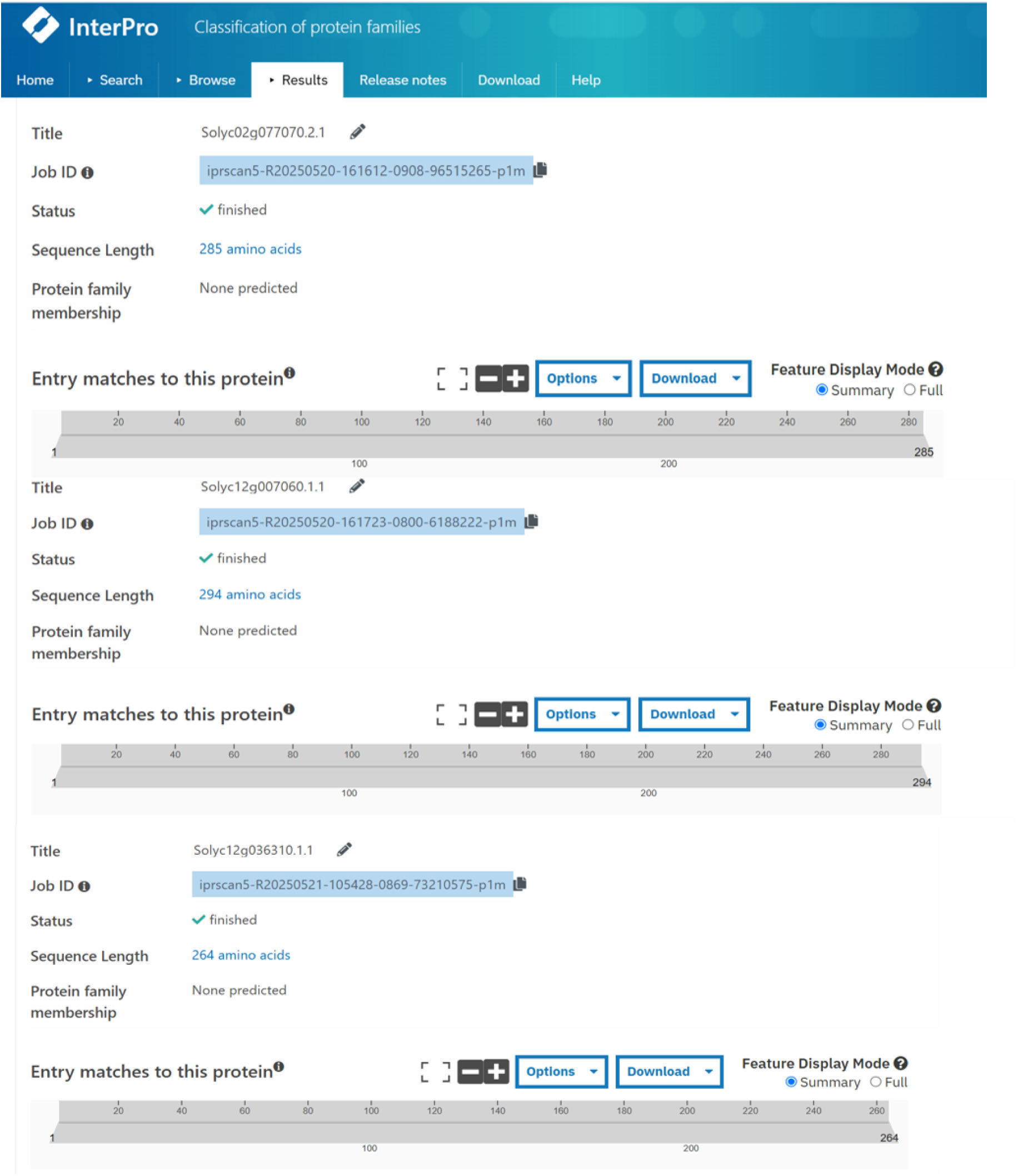
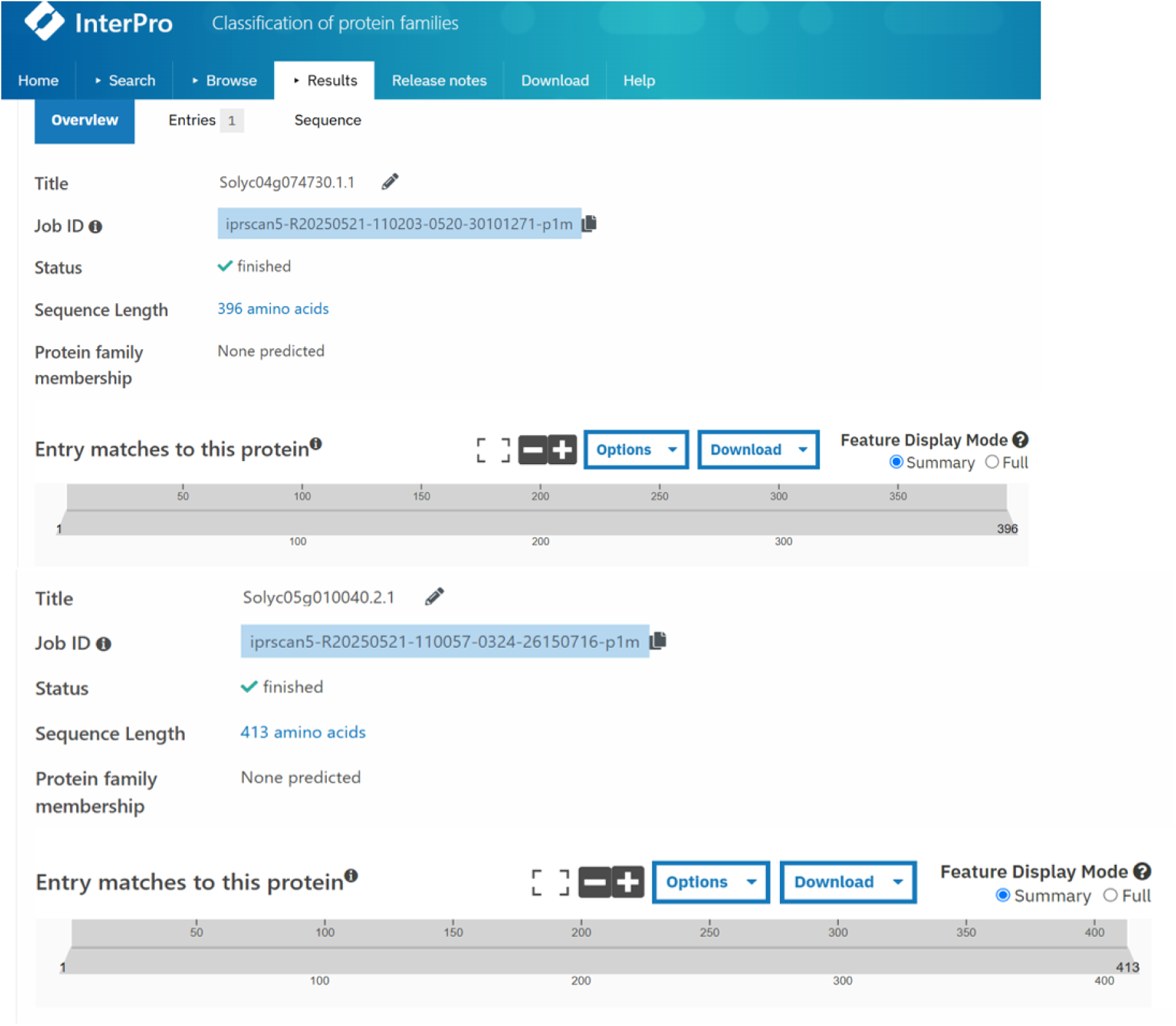
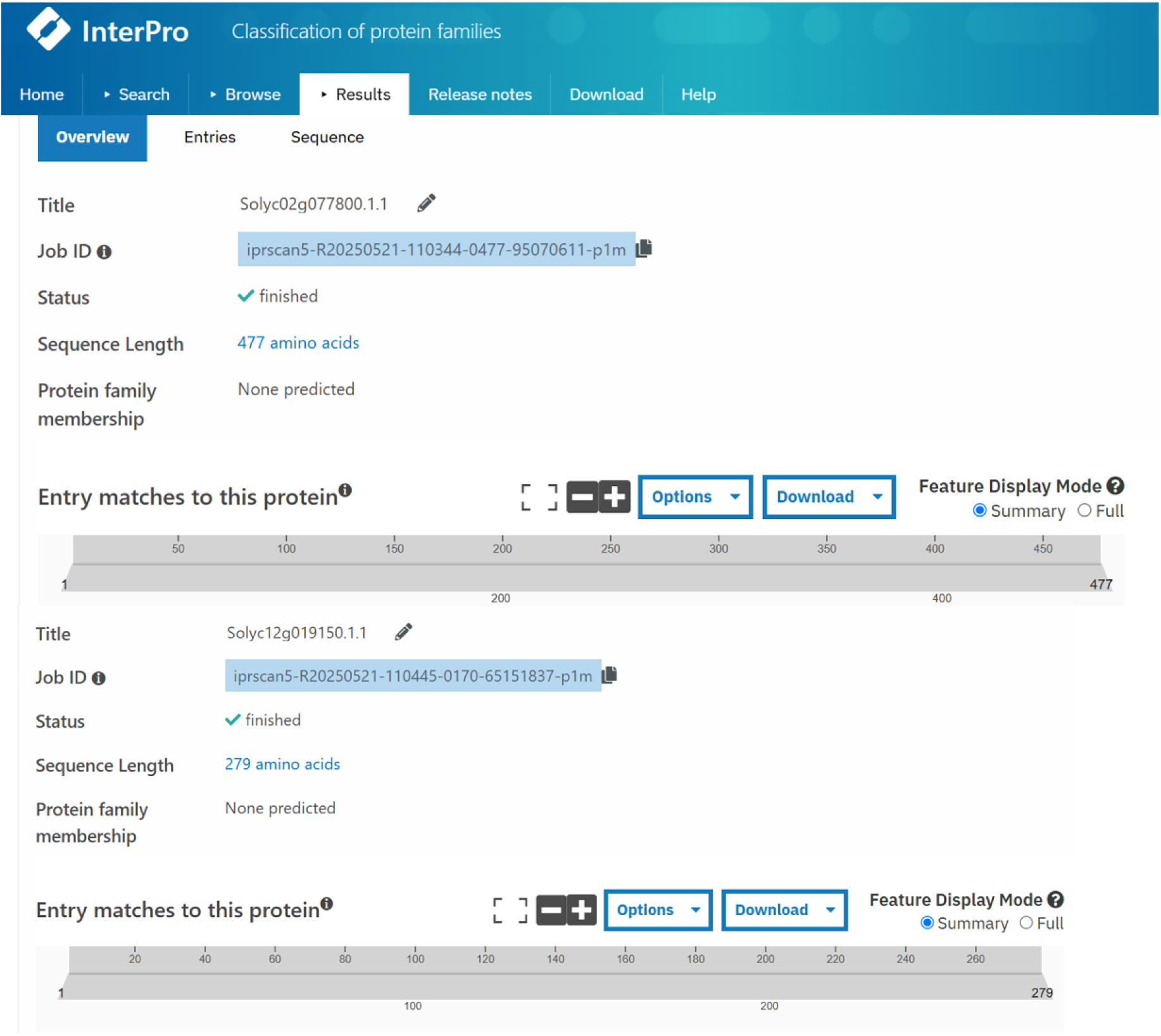
Potential LncRNA sequences searched against the InterPro database depicting the absence of protein-coding families.

## Notes

### Competing Interest Statement

The authors have declared no competing interest.

